# Homeostatic feedback between lysosomal mTORC1 and mTORC2-AKT signaling controls nutrient uptake in brown adipose tissue

**DOI:** 10.1101/2022.05.17.492242

**Authors:** Gudrun Liebscher, Nemanja Vujic, Renate Schreiber, Markus Heine, Caroline Krebiehl, Madalina Duta-Mare, Giorgia Lamberti, Cedric H. de Smet, Michael W. Hess, Thomas O. Eichmann, Sarah Hölzl, Ludger Scheja, Joerg Heeren, Dagmar Kratky, Lukas A. Huber

## Abstract

In brown adipose tissue (iBAT), the balance of lipid/glucose uptake and lipolysis is regulated by insulin signaling. Downstream of the insulin receptor, PDK1 and mTORC2 phosphorylate AKT, which activates glucose uptake and lysosomal mTORC1 signaling. The latter requires the late endosomal/lysosomal adaptor and MAPK and mTOR activator (LAMTOR/Ragulator). Deletion of LAMTOR2 (and thereby loss of the LAMTOR complex) in mouse adipocytes resulted in insulin-independent AKT hyperphosphorylation in iBAT, causing increased glucose and fatty acid uptake as evidenced by massively enlarged lipid droplets. As LAMTOR2 was essential for the upregulation of *de novo* lipogenesis, LAMTOR2 deficiency triggered exogenous glucose storage as glycogen in iBAT. These effects are cell autonomous, since AKT hyperphosphorylation was reversed by PI3K inhibition or by deletion of the mTORC2 component Rictor in LAMTOR2-deficient mouse embryonic fibroblasts. We identified a homeostatic circuit connecting LAMTOR-mTORC1 signaling with PI3K-mTORC2-AKT signaling downstream of the insulin receptor to maintain iBAT metabolism.

## Introduction

Adaption to the environment is an essential function of life that is executed by cells in response to e.g. nutrient availability and temperature changes. In mammals, cold exposure stimulates the secretion of norepinephrine from the sympathetic nervous system, which activates β_3_ adrenergic receptors at the surface of brown adipocytes. Consequently, non-shivering thermogenesis is induced to produce heat for maintaining body temperature^1^. This process requires a high amount of fatty acids (FA) and glucose, which are used as fuel and combusted in the mitochondria to build the proton gradient in the inner mitochondrial membrane^2^. The proton gradient is used for heat generation by the uncoupling protein 1 (UCP1)^3, 4^. To maintain supply for high energy needs, brown adipose tissue (iBAT) utilizes nutrient (FA and glucose) uptake as well as mobilization of FA from endogenous triglycerides (TG) by lipolysis^2^. Nevertheless, it has been suggested that in brown adipocytes lipolysis is not essential for non-shivering thermogenesis^5, 6^, arguing in favor of potentiated nutrient uptake in iBAT. Upon cold exposure and lipoprotein lipase (LPL)-mediated hydrolysis of TG-rich lipoproteins (TRL), brown adipocytes take up released FA by the FA transporter cluster of differentiation 36 (CD36)^7, 8^.

Recently, insulin signaling during cold stimulation has been reported to increase FA and glucose uptake from the blood^9^. Binding of insulin to the insulin receptor leads to the phosphorylation of the insulin receptor substrate (IRS) and to the activation of phosphoinositide-3-kinase (PI3K). PI3K, together with phosphatase and tensin homolog deleted on chromosome 10 (PTEN), regulates the abundance of phosphatidylinositol-3,4,5-trisphosphate (PIP3) in the plasma membrane and thereby allows the binding of phosphoinositide-dependent kinase 1 (PDK1). Subsequently, protein kinase B (PKB/AKT) is recruited to these domains and is phosphorylated by PDK1 at T308 and by the mechanistic target of rapamycin complex 2 (mTORC2) at S473^10–12^. Downstream of AKT, phosphorylation of the GTPase activating protein AKT substrate of 160 kDa (AS160, also known as TBC1D4) leads to the translocation of Solute carrier family 2, facilitated glucose transporter member 4 (GLUT4)-positive vesicles to the plasma membrane and allows a high influx of glucose into the cell^13^. AKT activation also mediates the activation of mTORC1 signaling at lysosomes by phosphorylating the RHEB inhibitor tuberculous sclerosis complex (TSC)^14^. For mTORC1 activation, the complex needs to be recruited to the surface of lysosomes. This is orchestrated by the late endosomal/lysosomal MAPK and mTOR activator (LAMTOR/Ragulator) complex and small Rag GTPases in the presence of amino acids. Subsequently, RHEB, located at the lysosomal surface, activates lysosomal mTORC1^15–17^.

The role of mTORC1 signaling in adipose tissue physiology has been demonstrated in several studies^11, 18–20^. In mouse iBAT, activation of mTORC1 affects non-shivering thermogenesis via activation of glucose and FA oxidation through the citric acid cycle^18, 21^. Furthermore, it regulates *de novo* lipogenesis by phosphorylation (and thereby inactivation) of lipin1 to promote the nuclear translocation of sterol regulatory element-binding protein 1c (SREBP1c)^22^. Although mTORC1 is not sufficient to induce SREBP1c, AKT is essential for insulin stimulated SREBP1c activation^23^. Besides being an integral part of mTORC1 signaling necessary for its lysosomal activation, LAMTOR also provides a scaffold for 5’- AMP-activated protein kinase (AMPK) and mitogen-activated protein kinase (MAPK) signaling^24–26^. From this perspective, LAMTOR integrates different anabolic as well as catabolic signals to balance metabolic signaling pathways. As they are all involved in the process of iBAT activation and regulation, we studied the effects of adipocyte specific LAMTOR deficiency on adipose tissue homeostasis in mice.

Here, we show that a feedback loop between LAMTOR-mediated lysosomal mTORC1 and mTORC2 signaling at the plasma membrane orchestrates the regulation of energy uptake and storage in iBAT. Mice lacking LAMTOR2, and thus the entire LAMTOR complex^27^, in adipocytes did not exhibit an obvious phenotype in the gonadal white adipose tissue (gWAT), but accumulated TG and glycogen in their iBAT and over time TG in the liver. In brown adipocytes, LAMTOR2 deficiency caused AKT hyperphosphorylation, which resulted in higher glucose and FA uptake. Loss of LAMTOR2 in mouse embryonic fibroblasts (MEF) recapitulated the phenotype and caused a similar hyperphosphorylation of AKT at S473, which was abolished by PI3K inhibition and LAMTOR2 / Rictor double deficiency. Overall, our study provides evidence that the LAMTOR complex is necessary to switch off AKT phosphorylation downstream of the insulin receptor, independently of previously established negative feedback loops^28–32^.

## Results

### Adipocyte-specific LAMTOR2-deficient mice accumulate lipids in their iBAT

To investigate the role of the LAMTOR complex in adipocytes, we generated mice specifically lacking LAMTOR2 (LT2) in white and brown adipocytes using the Adipoq-Cre driven loxP-system (LT2 AKO). Efficient gene deletion was confirmed by its reduced mRNA and protein expression in iBAT, inguinal WAT (iWAT), and gWAT, but not in the liver (Fig 1A, B). Loss of one LAMTOR subunit causes complex destabilization and degradation of all other four subunits, consequently leading to the breakdown and degradation of the LAMTOR complex as a whole^27^. In line, LAMTOR1 (LT1) protein levels were diminished in iBAT and gWAT of LT2 AKO mice (Fig 1B). Thus, LT2 AKO mice lacked the entire LAMTOR complex in adipocytes.

**Figure 1.**
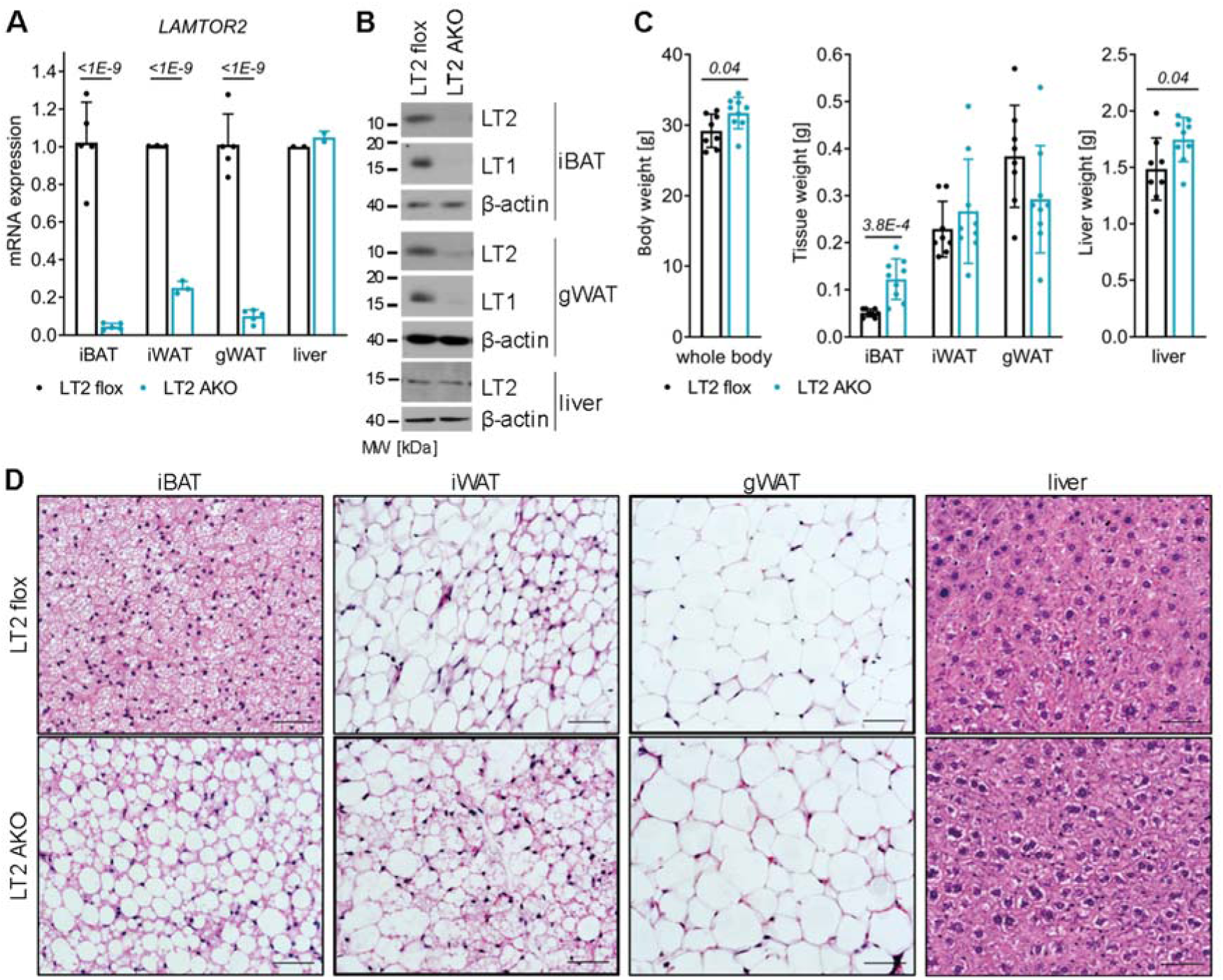
Pronounced lipid accumulation in iBAT of LT2 AKO mice. Male LT2 flox and LT2 AKO mice aged 12 weeks were assayed for (A) relative *LAMTOR2* mRNA expression in iBAT, iWAT, gWAT, and liver by qRT-PCR (n = 5) and (B) LAMTOR2 protein expression in lysates of iBAT, gWAT, and liver by Western blotting (n = 3). (C) Body weight and weight of iBAT, iWAT, gWAT, and liver (n=8). (D) Representative H&E staining of iBAT, iWAT, gWAT, and liver sections of mice housed at room temperature (scale bar 50 µm). Bar charts present means ± SD with individual values presented as dots. Statistical significance was calculated using two-tailed unpaired Student’s t-tests with italic numbers representing the p-value.

LT2 AKO animals exhibited an 8% increase in body weight (Fig 1C). Tissue analysis revealed a 2-fold increase in iBAT weight, and although iWAT and gWAT weights were unaffected, the liver weight was increased by 17% (Fig 1C). The iBAT weight gain was associated with a substantial accumulation of lipids and resulted in large, unilocular lipid droplets (LD) in LT2 AKO mice compared to the small multilocular LD in LT2 flox controls (Fig 1D). While gWAT of LT2 AKO mice was unaffected, in iWAT areas of adipocytes containing a large amount of smaller LD were evident (Fig 1D).

We observed that the iBAT and liver phenotype persisted in LT2 AKO mice maintained under thermoneutral conditions (28-30°C) for 5 weeks when BAT was not activated, as iBAT weight and LD size were also increased (Fig S1A-C). These data suggest that the lipid accumulation is independent of the thermoregulatory circuit. As loss of LT2 had the most striking effect on iBAT morphology, we further focused on the role of the LAMTOR complex in iBAT metabolism and the underlying mechanisms for the observed lipid accumulation.

The WAT-like phenotype of the iBAT in LT2 AKO mice led us to address if and how the iBAT function was regulated by lysosomal mTORC1 signaling. As a first characterization, we performed a microarray experiment to analyze the expression levels of brown and white adipocyte marker genes in iBAT using the ProFAT analysis tool^33^, which quantifies the thermogenic potential of adipose tissue samples. LT2 AKO iBAT expressed all BAT marker genes and the expression results clustered with the control iBAT samples (Fig. S1D, E), indicating classic iBAT functionality, despite the dramatic morphological changes and apparent whitening associated with loss of LT2. As control, gWAT samples of LT2 AKO and control mice similarly expressed the WAT marker genes and showed no expression of BAT marker genes (Fig. S1D, E).

Consistent with normal brown adipocyte gene expression, LT2 AKO mice were able to maintain their body temperature when housed at 5°C for 48h similar to LT2 flox control mice (Fig 2A). They upregulated upon cold exposure in iBAT the thermogenesis marker UCP1, which showed high variation in iBAT homogenates of LT2 AKO mice housed at room temperature (Fig 2B). Remarkably, after cold exposure for 48h, the LT2 AKO iBAT lipid accumulation and weight increase was no longer present (Fig 2C) and the body weight of LT2 AKO mice and their respective controls remained unchanged (Fig 2D). To evaluate the reversibility of the lipid accumulation in LT2 AKO iBAT, we housed the mice first at 5°C for 48h and subsequently at 22°C. Surprisingly, 24h at 22°C was sufficient for LT2 AKO iBAT to re-accumulate lipids (Fig 2E) while the body weight remained unchanged compared to LT2 flox control mice (Fig 2F). This result indicated that the iBAT of LT2 AKO mice accumulated lipids at 22°C but was capable to deplete these surplus lipids upon cold exposure.

**Figure 2.**
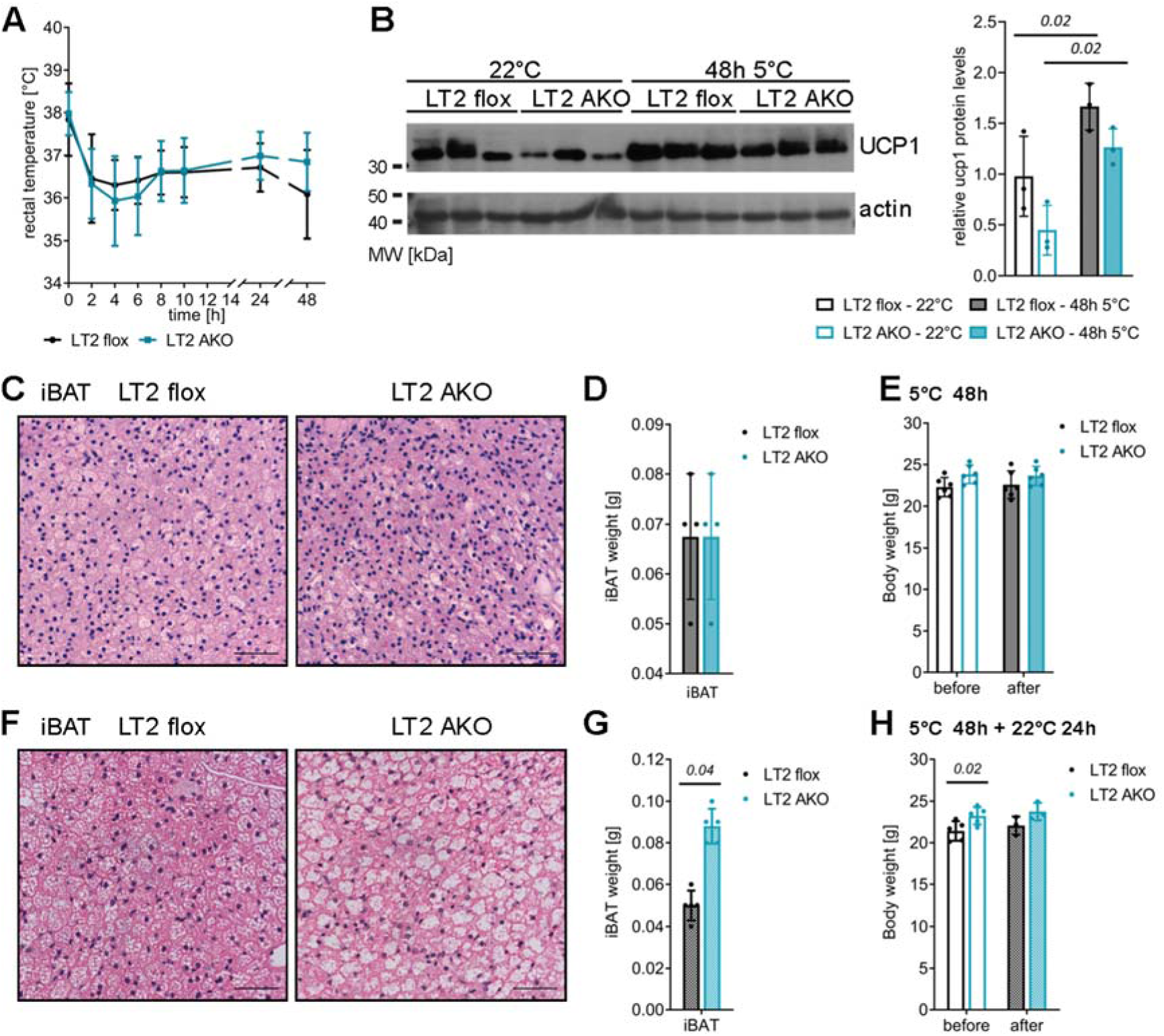
LT2 AKO mice are cold tolerant and deplete their excess iBAT lipids upon cold exposure, which re-accumulate within 24h at room temperature. (A) Rectal temperature of female mice housed for 48h at 5°C presented as means ± SD (n=7). (B) Immunoblots and quantification of UCP1 in iBAT lysates of male (22°C) and female mice (for 48h at 5°C) (n = 3). (C, F) Representative H&E staining of iBAT sections (scale bar 50 µm), (D, G) iBAT weight and (E, H) body weight of female mice housed for 48h at 5°C (C-E) without or (F-H) with subsequent 22°C housing for 24h (n=4). Bar charts present means ± SD with individual values presented as dots. Statistical significance was calculated using two-tailed unpaired Student’s t-tests with italic numbers representing the p-value.

### Reduced mTORC1 signaling, but AKT hyperphosphorylation in iBAT of LT2 AKO mice

Since the LAMTOR complex has a key role in lysosomal mTORC1 signaling, we investigated mTORC1 activation and the negative feedback loop to the insulin signaling pathway^34^ in iBAT. In protein extracts of iBAT, the total protein levels of Raptor and Rictor, the specific regulatory subunits of mTORC1 and mTORC2, respectively, were comparable between LT2 flox and AKO iBAT (Fig 3A). As refeeding causes activation of mTORC1 signaling^35–37^, mice were metabolically synchronized by fasting/refeeding. In the iBAT of refed LT2 AKO mice, the phosphorylation status of the mTORC1 downstream targets 40S ribosomal protein S6 (S6) and eukaryotic translation initiation factor 4E binding protein (4E-BP1) was markedly reduced (Fig 3B). In addition, mTORC1 phosphorylates the transcription factor EB (TFEB) and subsequently reduces the expression of its downstream targets^38, 39^. One of the most regulated TFEB targets is Ras-related GTP-binding protein D (RagD)^40^. Increased mRNA levels of *RagD*, *mucolipin1* (*MCOLN1*), *CD63* and *cathepsin a (CTSa*) in LT2 AKO iBAT (Fig 3C) indicated increased transcription of these known TFEB targets, due to reduced mTORC1 activity. In line with a recent publication investigating TFEB deficiency^41^, microarray analysis failed to reveal a systemic mRNA upregulation of the coordinated lysosomal expression and regulation (CLEAR) network. Nevertheless, some CLEAR genes such as *CD63*, *beta-hexosaminidase subunit β*, *MCOLN1*, *CTSa,* and *CTSz* were significantly upregulated in the microarray analysis (data not shown). In conclusion, loss of LT2 led to reduced LAMTOR-mediated lysosomal mTORC1 activity in iBAT.

**Figure 3.**
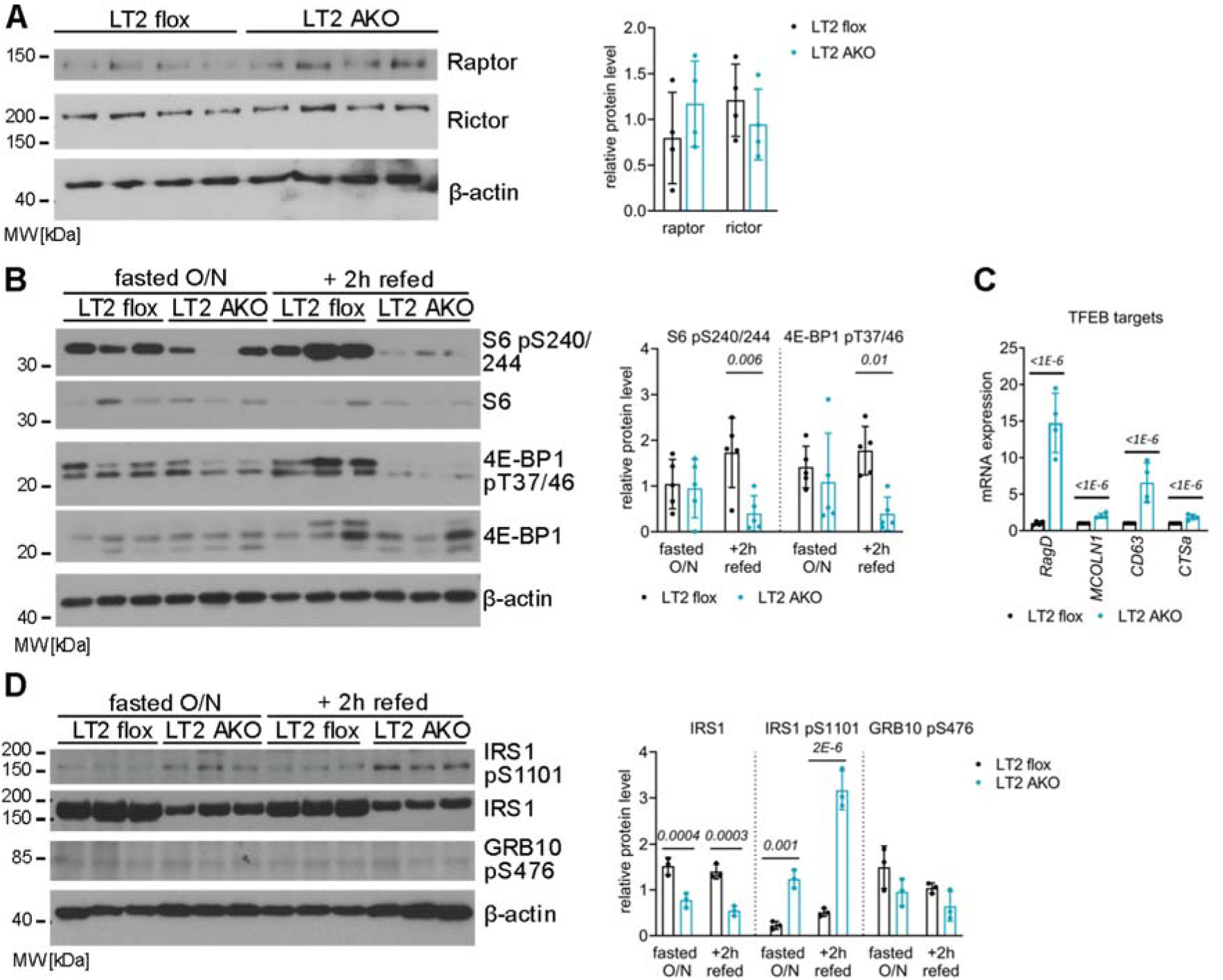
LAMTOR deletion in iBAT impairs mTOR signaling without affecting the negative feedback loop. Male LT2 flox and LT2 AKO mice were housed at 22°C. (A) Immunoblots and quantification of Raptor and Rictor in iBAT lysates (n=4). (B) Immunoblots and quantification of the mTORC1 downstream targets S6 and 4E-BP1 and their phosphorylation sites (S6 pS240/244, 4E-BP1 pT37/46) (n=5). (C) Relative mRNA expression of the TFEB downstream targets *RagD, MCOLN1, CD63* and *CTSa* in iBAT (n=4). (D) Immunoblots and quantification of total IRS, IRS pS1101, and GRB10 pS475 in iBAT lysates of male mice fasted overnight with or without refeeding for 2h with chow diet (n=3). Statistical significance was calculated using two-tailed unpaired Student’s t-tests with italic numbers representing the p-value.

In comparison, the classical targets of mTORC1 – S6 and 4E-BP1 – were partially downregulated or not conclusive regulated in ingWAT and gWAT, respectively (Fig S2 A, B). These results show that these white adipocytes may not be regulated by lysosomal mTORC1 signaling as brown adipocytes are.

To maintain sensitization to insulin, mTORC1 inhibits insulin signaling via phosphorylation of S6 kinase (S6K) and growth factor receptor-bound protein 10 (GRB10), which subsequently phosphorylate insulin receptor substrate 1 (IRS1), causing IRS1 degradation^28–32^. In both overnight-fasted and refed mice, the total amount of IRS1 in LT2 AKO iBAT was reduced, whereas phosphorylation at S1101, which activates degradation, was increased (Fig 3D). However, GRB10 phosphorylation remained unchanged (Fig 3D). These findings indicated that the insulin receptor pathway was reduced in LT2 AKO iBAT by the degradation of IRS1 due to activation of the negative feedback loop from mTORC1 to IRS1.

The plasma membrane-enriched fractions of LT2 AKO iBAT also contained a (albeit not significant) lower amount of mature insulin receptor β (Fig 4A). Downstream of the insulin receptor, mTORC2 and PDK phosphorylate and activate AKT at S473 and T308, respectively^10–12^. Of note, phosphorylation at both sites was strongly upregulated in iBAT of LT2 AKO mice that were either fasted overnight or refed for 2h (Fig 4B). Likewise, in iBAT of 4h-fasted LT2 AKO mice, we observed markedly enhanced phosphorylation of AKT (Fig. S3A). However, insulin injection increased AKT phosphorylation in LT2 flox iBAT to levels present in LT2 AKO iBAT (Fig S3A), indicating that insulin failed to further stimulate AKT phosphorylation in the iBAT of LT2 AKO mice. Thus, LT2 was required to facilitate insulin signaling in iBAT.In summary, LT2 was required to activate and stabilize IRS1, maintain the insulin receptor, and to attenuate mTORC2-dependent AKT activation in iBAT.

**Figure 4.**
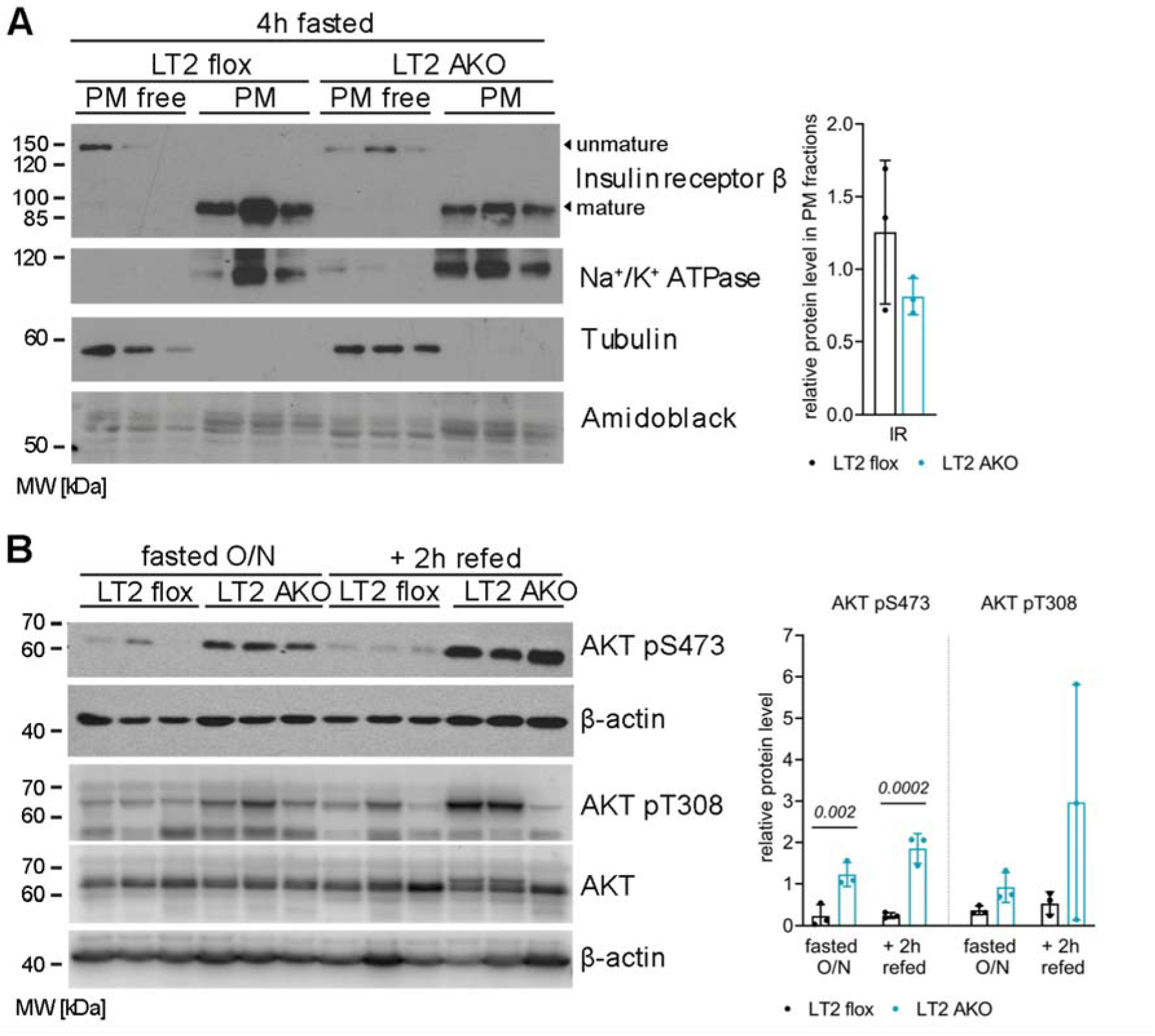
Reduced insulin receptor abundance at the plasma membrane, but hyperphosphorylated AKT. (A) Immunoblots and quantification of the insulin receptor, Na+/K+ ATPase, tubulin, and amidoblack staining of plasma membrane-free and -enriched fractions of iBAT lysates of 4h-fasted male mice housed at 22°C (n=3). (B) Immunoblots and quantification of AKT together with its respective phosphorylation sites (AKT pS475, AKT pT308) in iBAT lysates of overnight fasted male mice with or without refeeding with chow diet for 2h (n=3). Statistical significance was calculated using two-tailed unpaired Student’s t-tests with the italic numbers representing the p- value.

In LT2 AKO ingWAT a similar AKT hyperphosphorylation was observed (Fig S3B) but was lacking in LT2 AKO gWAT (Fig S3C). In conclusion, the role of the LAMTOR complex differ when comparing the different adipose tissue depots.

### LT2 affects AKT phosphorylation via PI3K and/or mTORC2

Next, we addressed whether LT2-dependent regulation of mTORC2 signaling was cell autonomous and/or iBAT specific. PI3K and mTORC2 are responsible for full AKT activation. Thus, we analyzed the cell autonomous regulation of these kinases in LT2-/- MEFs in comparison to LT2f/- and HA-LT2 reconstituted (LT2-HA) MEFs. We cultured these cells in serum-free medium (w/o FBS) and determined the phosphorylation of S6 and the consequences of PI3K inhibition by using ZSTK-474^42^. Consistent with a reduction in mTORC1 signaling, phosphorylated S6 was very low in LT2-/- compared to LT2 fl/- and to the reconstituted LT2-HA MEFs under all conditions investigated (serum starvation, insulin treatment) (Fig 5A, S4A). In contrast, the phosphorylation of AKT by mTORC2 at S473 under steady state, starvation, and insulin stimulation was increased only in LT2-/- MEFs (Fig 5A, S4B). A similar but weaker effect was observed for the PDK1-dependent phosphorylation of AKT at T308 (Fig 5A, S4C). Treatment of MEFs with the PI3K inhibitor ZSTK-474 showed in the absence and presence of LT2 a pronounced reduction of AKT phosphorylation at S473 and T308 in combination with insulin stimulation (Fig 5A, S4A-C). These findings suggested that LAMTOR affected PI3K itself or its downstream effectors PDK1 and mTORC2. To elucidate whether mTORC2, the kinase phosphorylating AKT at S473, might be involved, we generated LT2-/- Rictor-/- MEFs. S6 phosphorylation was absent in both LT2-/- and LT2-/- Rictor-/- MEFs but recovered after reconstitution of LT2 (Fig 5B, S4D). Of note, LT2-/- Rictor-/- MEFs failed to show phosphorylation of AKT at S473, independent of the culturing conditions (serum starvation, insulin treatment) (Fig 5B, S4E), despite the presence of AKT phosphorylation at T308 (Fig 5B, S4F).

**Figure 5.**
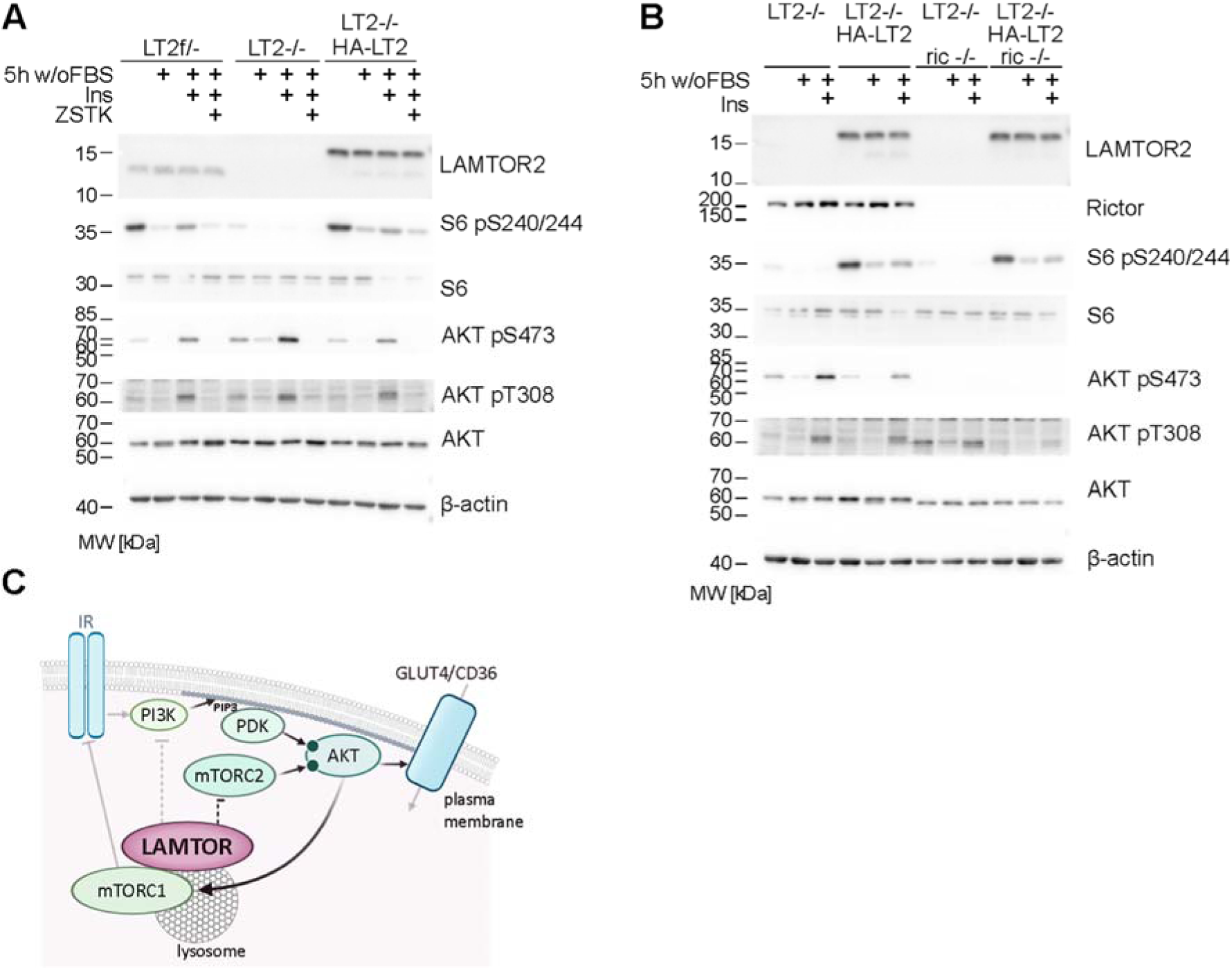
Inhibiting PI3K or deleting Rictor in LT2-/- MEFs reverses AKT hyperphosphorylation. LT2 fl/-, LT2-/-, LT2-/- with reconstituted (HA-)LT2, LT2-/- Rictor-/- and LT2-/- Rictor-/- HA-LT2 MEFs were harvested in steady state conditions (control) or after 5h of FBS starvation (w/o FBS) with or without PI3K inhibition (1 µM ZSTK-474 for 15 min) in the absence or presence of 100 nM insulin for 10 min. (A) Representative immunoblots of LT2, S6, and AKT together with their respective phosphorylated forms (S6 pS240/244, AKT pS475, AKT pT308) in lysates of LT2 fl/-, LT2 -/-, and LT2-/- HA-LT2 MEFs. For quantification see FigS3 (n=4). (B) Representative immunoblots of Rictor, LT2, S6, and AKT together with their respective phosphorylated forms (S6 pS240/244, AKT pS475, AKT pT308) in lysates of LT2-/-, LT2-/- HA-LT2, LT2-/- Rictor-/-, and LT2-/- Rictor-/- HA-LT2 MEFs. For quantification see FigS3 (n=4). (C) The LAMTOR complex is essential to switch off the signaling cascade downstream of the insulin receptor. The LAMTOR complex in iBAT positively regulates lysosomal mTORC1 activity and negatively regulates mTORC2 and/or PI3K activity toward AKT.

Hence, inhibition of PI3K or loss of mTORC2 signaling reversed the increased phosphorylation of AKT at S473 of LT2-/- MEFs. These findings suggested a regulatory circuit of lysosomal LAMTOR to PI3K and TORC2 at the plasma membrane to shut down insulin-AKT signaling (Fig 5C).

### Highly active AKT in LT2 AKO iBAT is associated with increased glucose and FA uptake

To evaluate the functional consequences of AKT hyperphosphorylation in LT2 AKO mice, we examined whether AKT downstream effectors could explain lipid accumulation in iBAT. In principle, increased FA uptake, increased glucose uptake and subsequent *de novo* lipogenesis, and decreased lipolysis could explain the phenotype. Due to the rapid lipid re-accumulation of lipids in iBAT after cold exposure, we focused on FA and glucose uptake. The abundance of the fatty acid transporter CD36 at the plasma membrane is increased by insulin^43^. AKT activation promotes cell surface accumulation of GLUT4 through phosphorylation of the Rab7 GTPase-activating protein AS160, which facilitates translocation of GLUT4-containing vesicles to the plasma membrane^44–46^. Indeed, the hyperactivation of AKT in LT2 AKO iBAT resulted in increased phosphorylation and thus inhibition of AS160 in both fasted and refed states (Fig 6A). Consistently, the significantly higher amounts of GLUT4 and CD36 were detected in plasma membrane fractions from iBAT of 4h-fasted LT2 AKO mice (Fig 6B).

**Figure 6.**
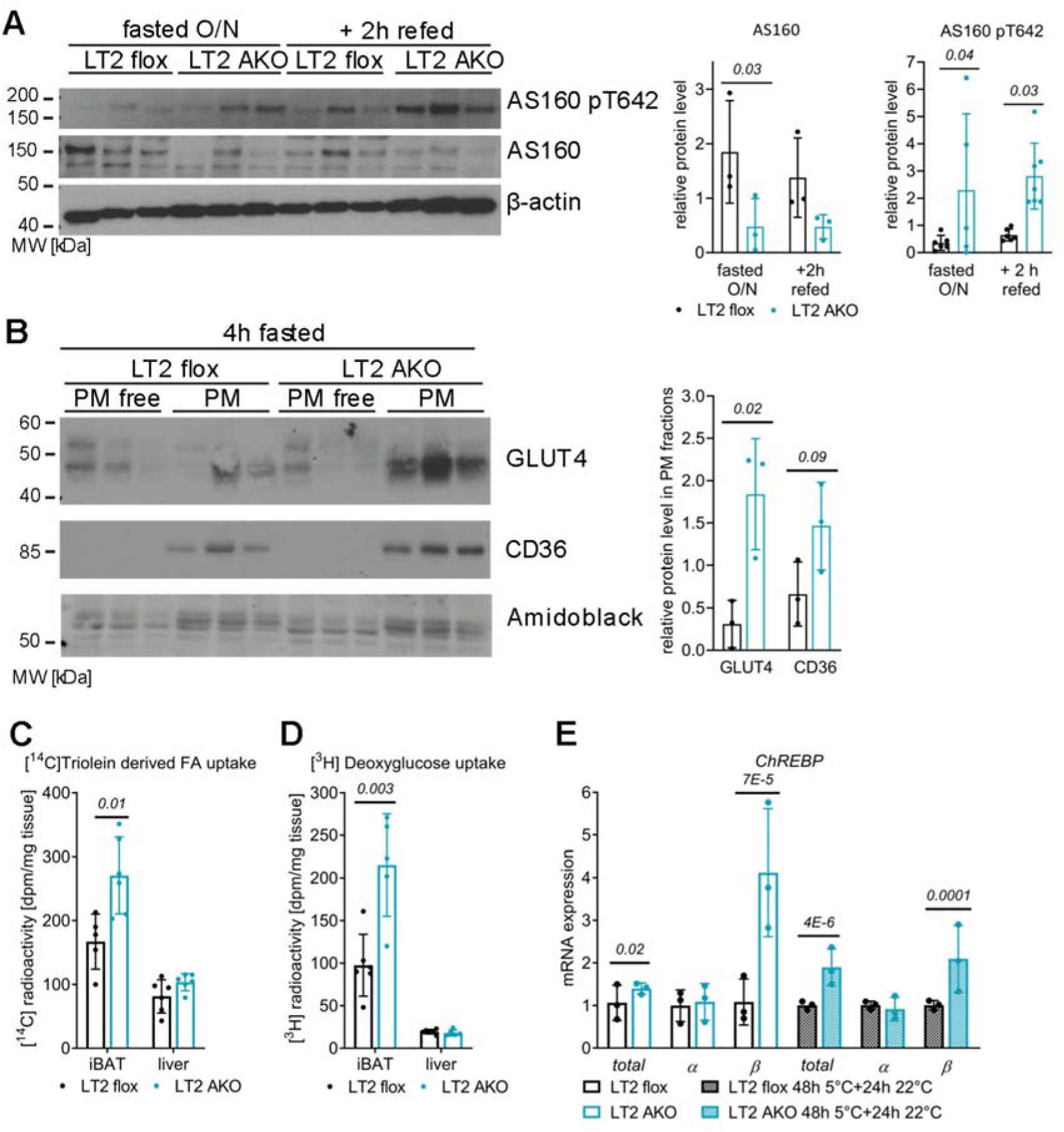
Hyperphosphorylated AKT in LT2 AKO iBAT is associated with increased CD36 and GLUT4 abundance at the plasma membrane and glucose and FA uptake. (A) Immunoblots and quantification of AS160 and AS160 pT642 in iBAT lysates of overnight fasted male mice with or without refeeding with chow diet for 2h (n=5). (B) Immunoblots and quantification of GLUT4, CD36, and amidoblack staining (same as in Fig.4 as same membrane) of plasma membrane-free and -enriched fractions of iBAT lysates of 4h-fasted male mice housed at 22°C (n=3). Male mice were fasted for 4h before they were (C) gavaged with intralipid emulsion containing [^14^C]triolein and (D) i.p. injected with [^3^H]deoxyglucose and uptake of radioactivity was determined in iBAT and liver (n=6). (E) Relative mRNA expression of total *ChREBP*, *ChREBPα*, and *ChREBPβ* in iBAT of male mice housed at 22°C or female mice housed at 5°C for 48h and afterwards at 22°C for 24h (n=3). Bar charts present means ± SD with individual values presented as dots. Statistical significance was calculated using two-tailed unpaired Student’s t-tests with italic numbers representing the p-value.

In line with this observation, the uptake of orally administered [^14^C]triolein-derived FA and of i.p. injected [^3^H]deoxyglucose was increased by 1.6- and 2.2-fold, respectively, in iBAT but not in livers of LT2 AKO mice (Fig 6C, D). Similarly, in LT2 AKO ingWAT and gWAT glucose uptake was increased by 3- and 2- fold, respectively (Fig S6A, C). But only in LT2 AKO ingWAT the FA uptake was additionally increased by 1.8-fold (Fig S6B, D). This enhanced glucose and FA uptake is not a consequence of changes in plasma insulin or blood glucose levels, LPL protein expression, or plasma lipid concentrations (Fig S6E- G). Although glucose tolerance was unaffected (Fig S6H), mTORC1-mediated inhibition of IRS may have caused the observed insulin resistance in LT2 AKO mice (Fig S6I). However, the respiratory exchange ratio and energy expenditure were unaltered in LT2 AKO mice housed either at room temperature or at 5°C (Fig S7A>, B), suggesting that LAMTOR specifically regulates substrate uptake and utilization but not energy combustion.

Consistent with increased glucose uptake, mRNA expression of carbohydrate-responsive element- binding protein β (*ChREBPβ*) was 3.8- and 2.1-fold higher in LT2 AKO mice housed at 22°C or at 5°C for 48h with subsequent 24h at 22°C, respectively (Fig 6E). Hence, LAMTOR mediated lysosomal mTORC1 restricted glucose and FA uptake in iBAT by controlling mTORC2-AKT signaling.

### LAMTOR2 deficiency dampens *de novo* lipogenesis activation and causes glycogen storage

The increased *ChREBPβ* expression along with elevated glucose uptake suggested increased *de novo* lipogenesis^47–49^. However, several genes involved in *de novo* lipogenesis were downregulated in LT2 AKO iBAT (*FASN* by 37%, *SCD1* by 62% and *ELOVL6* by 26%) (Fig 7A) as were FASN protein levels (Fig 7B). ChREBP acts together with SREBP1c as a transcription factor to activate genes involved in *de novo* lipogenesis^49–51^. Insulin and mTORC1 signaling trigger SREBP1c ER export, proteolytic processing, and entry of the N-terminal fragment into the nucleus to stimulate target gene transcription^22, 52^. Consistent with reduced mTORC1 signaling in LT2 AKO iBAT, cleaved and uncleaved SREBP1c did not increase upon refeeding in LT2 AKO iBAT, as observed in LT2 flox mice, but rather decreased (Fig 7C). SREBP1c levels remained similar in LT2 AKO iBAT among all treatments. An additional cause for decreased cleaved SREBP1c could be the increased free cholesterol content in LT2 AKO iBAT (Fig 7D), because high concentrations of free cholesterol inhibit SREBP1c proteolytic activation^53^. These results led us to predict that *de novo* lipid biosynthesis required LT2 and lysosomal mTORC1 signaling. In agreement with this prediction, [^14^C]glucose incorporation into TG and FA of LT2 AKO iBAT explants after cold exposure was reduced by 64% and 73%, respectively (Fig 7E). Instead of using the excess glucose for lipid biosynthesis, glucose accumulated as glycogen in LT2 AKO iBAT, as demonstrated by periodic acid Schiff (PAS) cytochemistry (Fig 7F) and glycogen quantification (Fig 7G). This correlated with the up-regulation of several genes involved in glycogenesis in iBAT of L2T AKO mice (Fig 7H).

**Figure 7.**
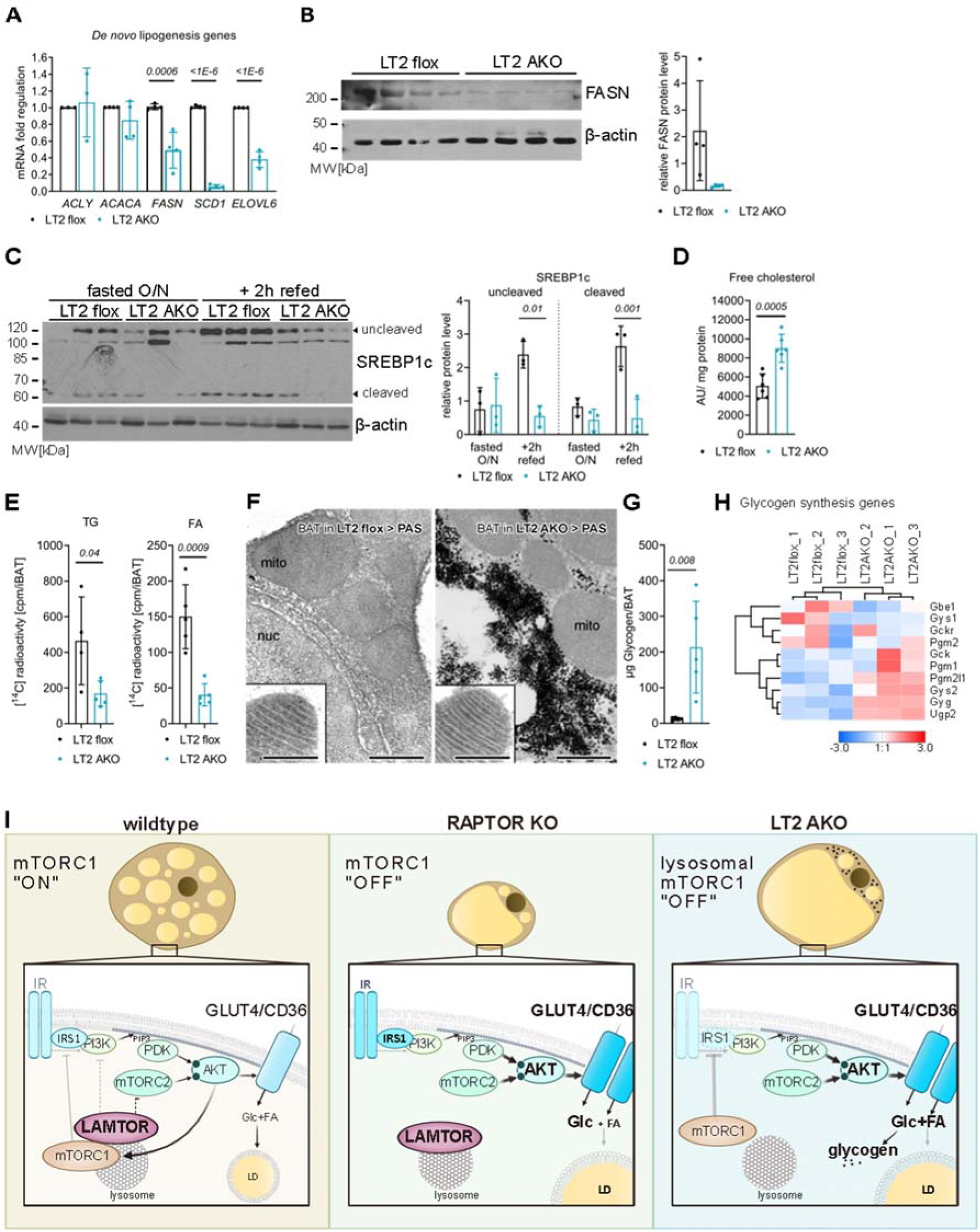
Reduced *de novo* lipogenesis in LT2 AKO iBAT shifts glucose to glycogen storage. Male LT2 flox and LT2 AKO mice were housed at 22°C. (A) Relative mRNA expression of the *de novo* lipogenesis genes *ACLY, ACACA, FASN, SCD1*, and *ELOVL6* in iBAT (n=3). (B) Immunoblots and quantification of FASN in iBAT lysates (n=4). (C) Immunoblots and quantification of SREBP1c in iBAT lysates of mice fasted overnight with or without refeeding for 2h with chow diet (n=3). (D) Quantification of free cholesterol levels in iBAT of 4h-fasted mice. (E) Quantification of [^3^H]glucose incorporation into iBAT explants of mice after they were exposed to 5°C for 48h (n=5). TG and FA were separated by TLC and radioactivity in the lipid bands was quantified by liquid scintillation counting. (F) Electron micrographs of iBAT resin sections subjected to PAS cytochemistry (Thiéry reaction) show glycogen accumulation (black reaction product) in LT2 AKO mice (mito, mitochondrion; nuc, nucleus). Inserts show identical, intact ultrastructure of mitochondria, as seen in cryo-sections (scale bar 500 nm). (G) Quantification of iBAT glycogen concentrations (n=5). (H) Clustered heatmap of microarray data of iBAT mRNA showing glycogen biogenesis associated genes (n=3). (I) A Raptor KO lacks mTORC1 and loses thereby the capability of IRS1 inhibition, which causes increased AKT phosphorylation and subsequent glucose uptake. Similarly, LAMTOR complex deficiency caused increased AKT phosphorylation, which increased GLUT4 and CD36 abundancy at the plasma membrane and thereby glucose and FA influx. In contrast, Glucose was mainly stored in form of glycogen, as *de novo* lipogenesis was not induced. Additionally, the mTORC1 complex was still partially active and able to inhibit IRS-1. As mTORC1 signalling is essential for cell growth, a lack of total mTORC1 reduces the cell size. But a lack in only lysosomal mTORC1 signalling has no effect on cell growth. Bar charts present means ± SD with individual values presented as dots. Statistical significance was calculated using two-tailed unpaired Student’s t-tests with italic numbers representing the p-value.

We concluded that the LAMTOR complex uses a homeostatic feedback mechanism to turn off mTORC2-AKT. While the exact mechanistic details remain to be identified, this feedback is essential to adapt FA and glucose uptake in the brown adipocytes and to funnel glucose into lipid biosynthesis (Fig 7I).

## Discussion

Signaling pathways are tightly regulated to allow fast response and adaption to incoming signals. Therefore, they must be turned off after major downstream targets have been reached. This is also true for the regulation of insulin signaling, which has been extensively studied in liver, skeletal muscle, and WAT^12, 54^. However, its role in BAT homeostasis, although important for nutrient uptake^9^, is less well understood.

Here, we describe the role of the LAMTOR complex in attenuating mTORC2-AKT signaling in iBAT. Our results open the exciting possibility that LAMTOR mediated lysosomal mTORC1 signaling feeds back to the plasma membrane to regulate AKT phosphorylation. However, the underlying molecular mechanism is not yet clear although preliminary data from our group suggest involvement of lysosomal localization (data not shown).

One of the major functions of the LAMTOR complex is being a scaffold for mTORc1 and thereby essential for lysosomal mTORC1 activation^17^. Hence, at first glance it doesn’t seem surprising that a LAMTOR KO shows AKT hyperphosphorylation comparable to mTORC1 inhibition e.g., by Adipoq-Cre-mediated Raptor deletion in adipose tissue^19^. Nevertheless, the observed phenotypes of Raptor or LAMTOR deletions correlate only partially^18, 19, 55^, such as insulin resistance in both mouse models^19^, and there are striking differences (Fig 7I). Despite a pronounced lipid accumulation in iBAT of Raptor- or LAMTOR-deficient mice, the iBAT weight is markedly reduced in Raptor AKO mice due to a proliferation defect^18^, which is absent upon LAMTOR deletion. Upon cold exposure, Raptor deficient mice are unable to increase the expression of *de novo* lipogenesis genes despite increased glucose uptake in iBAT^18^. This phenotype is similar to iBAT of LT2 AKO mice housed at room temperature. Although LT2 AKO iBAT exhibits increased glucose uptake *in vivo* and *ChREBPβ* gene expression, which is expected to potentiate *de novo* lipogenesis gene transcription^47^, glucose uptake in LT2 AKO iBAT explants resulted in reduced incorporation of glucose into FA and TG. Moreover, mRNA expression of *de novo* lipogenesis genes such as *FASN* and *SCD1* and its transcription factor SREBP1c are reduced. SREBP1c is probably regulated on at least two levels. First, the mTORC1 downstream target lipin-1 inhibits the nuclear entry of SREBP1c when mTORC1 is inactive^22^. Second, SREBP1c maturation might be inhibited by the increased free cholesterol content in LT2 AKO iBAT^56^. In addition, there might be an influence of AKT signaling on SREBP1c activation, which should be further investigated^23^.

Nevertheless, LT2 AKO mice tolerate cold exposure much better than Raptor deficient mice^18^. The reason for these differences could be a partially impaired activation of mTORC1 in LT2 AKO iBAT, as S6 and 4E-BP1 were still phosphorylated (to low levels), even under fasting conditions. This finding suggests residual mTORC1 activation independent of LAMTOR or at other organelles, which has been described previously in human and mouse cell models; e.g. LT2 deletion in dendritic cells^57^. However, even the inhibition of lysosomal mTORC1 activation profoundly influences iBAT homeostasis.

Surprisingly and unobserved in Raptor KO adipose tissue, LT2 AKO iBAT accumulates large amounts of glycogen. This suggests a shift from lipid to glucose metabolism upon LAMTOR deletion. Glycogen accumulation in iBAT was observed short time after refeeding^58^, after cold exposure^59^, and after disruption of TG synthesis by deletion of SCD1^60^ or DGAT2^61^ to store excess glucose. As LT2 AKO iBAT shows reduced *de novo* lipogenesis associated with increased glucose uptake, glycogen appears to be a logical way to store surplus glucose. In general, glycogen turnover plays an important role upon cold exposure, as it has been shown that it was necessary to induce e.g. UCP1 expression in iWAT and thus thermogenesis^62^. The importance and regulation of glycogen metabolism in BAT is poorly understood, but the high abundancy of glycogen synthesizing and metabolizing proteins^62^ suggest an important role that needs further investigation. Our data suggest that LAMTOR is important for glycogen homeostasis.

Raptor deficient iBAT showed a similar AKT hyperphosphorylation, which was explained as consequence of the negative feedback loop from mTORC1 to IRS-1 without investigating its activation^19^. In contrast in LT2 AKO mice, we observed a potentiated feedback mechanism through increased phosphorylation of IRS-1 and its degradation. This contradicts the hypothesis that reduced mTORC1 activation leads to a reduced negative feedback loop to the insulin receptor. Our results rather suggest the existence of another regulatory pathway, which is dysfunctional upon LAMTOR deletion. We hypothesize that the LAMTOR complex attenuates PI3K, mTORC2, and PDK1 activity, which explains the high AKT phosphorylation upon LAMTOR deletion.

Activation of mTORC2 and subsequently AKT was already shown in several publications to be important for insulin stimulated glucose uptake. Adipocyte specific Rictor knock-out (Rictor AKO) mice show reduced NEFA and TG uptake as well as reduced AKT2 dependent glucose uptake especially under cold stimulation^63^. Similarly, an myf5-lineage knock-out of Rictor shows reduced lipid accumulation of lipids in BAT due to a lipid metabolism defect, but a surprising increase in glucose uptake as they might switch main nutrient source^64^. This might be the result of a still partially phosphorylated AKT at Thr308. In addition, an adipocyte-specific AKT 1 and 2 knock-out mouse lacks subscapular white and brown adipose tissue depots^65^.

All these observations complicate the analysis which component of the insulin stimulated glucose uptake system is the key driver of the phenotype observed in LT2 AKO iBAT. Noteworthy, a LT2 KO might impact on AKT activation and the translocation of glucose sensitive vesicles to the plasma membrane due to an endosomal/lysosomal disorganization^66^. The delocalization of endosomes and lysosomes might interfere with the transport of GLUT4-containing vesicles to the plasma membrane. Furthermore, previous studies of the LAMTOR complex showed decreased endocytosis of receptors (epidermal growth factor receptor in keratinocytes and fms-like tyrosine kinase 3 ligand receptor in dendritic cells) in LAMTOR-deficient cells (Bohn et al., 2007; Scheffler et al., 2014). This might cause the accumulation of PIP3-enriched plasma membrane domains and enhance AKT phosphorylation at these domains. Some reports describe a varied importance of AKT isoforms in iBAT and insulin signaling. AKT-2 was proposed to be the major active isoform in iBAT. iBAT-specific AKT-2 deficiency mainly affected *ChREBPβ* expression and *de novo* lipogenesis, whereas other targets remained unchanged^50^. Interestingly, a study in hepatic cells suggested an activation of AKT-2 at PI(3,4)P2 domains of endosomes. This activation is hypothesized to allow prolonged AKT signaling to specific downstream targets^67, 68^. As a loss of LAMTOR changes the morphology and the dynamics in the endo-lysosomal system^66^, LAMTOR2 deletion might lead to a prolongation of AKT-2 activity at endosomes.

Similarly, delayed activation of AKT has been identified in biogenesis of lysosome-related organelles complex one-related complex (BORC)-deficient cells^69^. BORC is a lysosomal complex responsible for trafficking of lysosomes to the cell periphery. LAMTOR and BORC can interact with each other and this interaction regulates the positioning of lysosomes in response to signals such as amino acids and growth factors^70–72^. Furthermore, there might be little amounts of AKT and mTORC2 associated with lysosomes.^69^. As LAMTOR-deficient cells have an increased number of peripheral lysosomes^66^, these peripheral lysosomes could serve as an additional activation site for AKT due to their close proximity to the plasma membrane, PI3K, and PDK1. Preliminary data of our lab showed that moving lysosomes away from the periphery through a LT2-BORC double knock-out reverses AKT hyperphosphorylation of a LT2 single KO (data not shown). This indicates a regulation of AKT phosphorylation independent of mTORC1, but based on lysosomal positioning and abundancy. This is another aspect to distinguish a LT2 KO from mTORC1 inhibition and suggest a broader function of LT2. This hypothesis requires further investigation, particularly with regard to activation of the various AKT isoforms and lysosomal localization in a model system that is more suitable than brown adipocytes for spatially resolved microscopic studies. In conclusion, by using a deletion of LAMTOR in a highly insulin-responsive tissue, we discovered a new pathway for LAMTOR to change the insulin pathway from the lysosomal surface.

## Material and Methods

### Materials availability

All data generated or analyzed during this study are included in the manuscript and supporting file; source data files have been provided. The microarray dataset is available upon request.

### Animals

All mouse experiments were approved by the Austrian Federal Ministry of Science, Research, and Economy (BMBWF-66.011/0091-WF/II/3b/2014, BMBWF-66.011/0046-WF/V/3b/2015 and BMBWF-66.011/0006-V/3b/2019) and by the animal ethics advisory board of the Medical University of Innsbruck. The adipocyte-specific LAMTOR2 knockout mice (LT2 AKO) were generated by crossing LT2 flox/flox mice^73^ with Adiponectin-Cre: B6;FVB-Tg(Adipoq-cre)1Evdr/J mice^74^ (010803, Jackson Laboratory, Bar Harbor, ME). If not stated otherwise, age- and sex-matched mice aged between 12-16 weeks were used. The mice were maintained at standard housing conditions in a specific-pathogen-free (SPF) facility with a 12h light/12h dark cycle*, ad libitum* access to water and regular chow diet (V1534-300, ssnif, Soest, Germany) unless stated otherwise. For cold exposure, the mice were single-caged and the housing temperature was set to 5°C for 48h. The rectal temperature was measured using a rectal probe thermometer (Physitemp Instruments, Inc., Clifton, NJ). In addition, one cohort of mice was exposed to cold for 48h and then kept at room temperature for 24h. For the fasting/refeeding experiment, the mice were fasted overnight for 16h (16:00-08:00) and received afterwards chow diet for 2h (refeeding). For the thermoneutrality experiment, mice (8-9 weeks old) were kept for 5 weeks at 30°C with *ad libitum* access to water and regular chow diet. For the insulin treatment, mice were fasted for 4h and treated with 0.75 units insulin / kg body weight (INSUMAN # 1843315, Sanofi, Paris, France) for 15 min.

The mice were euthanized by cervical dislocation, the respective organs were harvested and snap-frozen in liquid nitrogen. For plasma analysis, the mice were fasted for 4h, blood was collected from the vena facialis, and plasma was prepared within 20 min. Plasma insulin, TG, NEFA, free glycerol, and cholesterol concentrations were measured using the respective kits according to manufacturers’ instructions (Mouse Ultrasensitive Insulin ELISA: 80-INSMSU-E01, Alpco, Salem, NH; Triglycerides Quantification Kit (183000), Cholesterol Quantification Kit (118000): Greiner Diagnostics, Bahlingen am Kaiserstuhl, Germany; Free Glycerol Reagent: F6428, Sigma-Aldrich, St. Louis, MO; NEFA Kit: WA434- 91795, WA436-91995, Wako Chemicals, Neuss, Germany). Blood glucose was measured by using an Accu-Check® Performa blood glucometer (Roche Diagnostics GmbH, Mannheim, Germany). Mice were excluded from the experiment when reaching a termination criterion such as hypothermia or a loss of 15% of their body weight.

All animal experimental experiments are described according to the ARRIVE guidelines.

### Histology

For hematoxylin and eosin staining (HE), iBAT, liver, iWAT, and gWAT were fixed for at least 24h in 4% buffered formaldehyde and embedded in paraffin. Thereafter, 5 µm thick sections were stained with HE using a standard protocol (Hematoxylin (3816.2), Eosin (0331.1): Carl-Roth, Karlsruhe, Germany). For Oil Red O staining (O0625, Sigma-Aldrich, St. Louis, MO) of cryo-sections, samples of liver were embedded in Tissue Tek O.C.T. (TTEK, A. Hartenstein, Würzburg, Germany), snap-frozen in liquid nitrogen, and cut into 5 µm thick cryo-sections fixed with 4% formaldehyde for 10 min.

### Gene expression analysis

RNA was isolated from iBAT, iWAT, gWAT, and liver using the RNeasy Kit (74104, Qiagen, Hilden, Germany). RNA integrity was validated by gel electrophoresis and 800 ng were reverse transcribed using the LunaScript® RT SuperMix Kit according to manufacturer’s instructions (E3010, New England Biolabs, Ipswich, MA). Real-time PCR was performed using the Blue ŚGreen qPCR Kit (F416, Biozym Scientific, Hessisch Oldendorf, Germany) with gene-specific primers (see Supplemetary Table 1) in duplicates or triplicates. Technical replicate outlier were excluded when determined by an outlier analysis. Gene expression was calculated using the 2^-ΔΔCt^ method and normalized to the housekeeper genes *TATA-box binding protein* (*Tbp*) and *40S ribosomal protein S20* (*Rps20*).

The Expression Profiling Unit of the Medical University of Innsbruck measured Affymetrix-based gene expression profiles as described previously^75^. Total RNA integrity was analyzed using the Bioanalyzer 2100 (Agilent Technologies, Vienna, Austria). Five hundred ng of high-quality RNA was hybridized to biotin and subsequently onto Affymetrix Mouse Genome 430 2.0 GeneChips according to the manufacturer’s protocols (900497, Applied Biosystems, Waltham, MA). Thereafter, the microarray was washed, stained in an Affymetrix fluidic station 450, and scanned in an Affymetrix scanner 3000. The results were analyzed using R version 3.0.3 (http://r-project.org) with the Bioconductor project package^76^. The data were preprocessed by the GCRMA method^77^. A moderated t-test was used for calculating the significance of the differential expression^78^ and the p-value was adjusted afterwards for multiple testing using the Benjamini and Hochberg method^79^. For pathway enrichment analysis, genes with a significant ≥1.4-fold up- or downregulation were analyzed by using the Enrichr webpage^80–82^ (http://amp.pharm.mssm.edu/Enrichr/). The results of the microarray analysis were used for a Prediction of adipocyte identity^33^ (ProFAT analysis: http://profat.genzentrum.lmu.de).

### Western blot analysis

For protein analysis, snap-frozen samples of iBAT, gWAT, and liver were homogenized in RIPA buffer (150 mM NaCl, 1.0% NP-40, 0.5% sodium deoxycholate, 0.1% SDS, 50 mM Tris, pH 8.0) or lysis buffer (0.25 M sucrose, 1 mM EDTA, 1 mM DTT, pH 7.0 with KOH) and freshly added protease and phosphatase inhibitors. The protein concentration was measured by using a BCA Protein Assay Kit (23225, ThermoFisher Scientific, Waltham, MA). Samples containing 20-50 µg protein were separated by SDS polyacrylamide gel electrophoresis and transferred to a nitrocellulose or polyvinylidene fluoride membrane (10600023, VWR, Radnor, PA). After blocking the membrane for 1h in 3% BSA in Tris-buffered saline with Tween 20 (TBS-T), the blots were probed with primary antibodies (Rabbit polyclonal anti-UCP1: ab10983, Abcam, Cambridge, United Kingdom; Rabbit monoclonal anti-LAMTOR1: HPA002997, Atlas Antibodies, Bromma, Sweden; Rabbit monoclonal anti-4E-BP1 (9644), Rabbit monoclonal anti-Phospho-4E-BP1 T37/46 (2855), Mouse monoclonal anti-Actin (clone C4) (3700), Rabbit polyclonal anti-AKT (9272), Rabbit polyclonal anti-Phospho-AKT S473 (4060), Rabbit polyclonal anti-Phospho-AKT T308 (9275, 13038), Rabbit monoclonal anti-AS160 (2670), Rabbit monoclonal anti-Phospho-AS160 T642 (8881), Rabbit polyclonal anti-Phospho-GRB10 S476 (11817), Rabbit monoclonal anti-Insulin receptor β (4B8) (3025), Rabbit polyclonal anti-IRS1 (3407), Rabbit polyclonal anti-Phospho-IRS1 S1101 (2385), Rabbit monoclonal anti-LAMTOR2 (8145), Mouse monoclonal anti- S6 Ribosomal Protein (2317), Rabbit polyclonal anti-Phospho-S6 Ribosomal Protein S240/244 (2215), Rabbit polyclonal anti-Raptor (2280), Rabbit monoclonal anti-Rictor (2114): Cell Signaling Technology, Danvers, MA; Mouse monoclonal anti-Tubulin: 12G10, Developmental Studies Hybridoma Bank, Iowa City, IA; Rabbit monoclonal anti-Na+/K+-ATPase: GTX61640, GeneTex, Irvine, CA; Mouse monoclonal anti-Actin (clone C4): MAB1501, Merck-Millipore, Darmstadt, Germany; Rabbit polyclonal anti-CD36 (NB400-144), Rabbit polyclonal anti-FASN (NB400-114SS): Novus Biologicals, Littleton, CO; Rabbit polyclonal anti-GLUT4: 21048-1-AP, ProteinTech Group, Manchester, United Kingdom; Mouse monoclonal anti-SREBP1c (Clone 2A4): sc-13551, Santa Cruz Biotechnology, Dallas, TX) overnight at 4°C. Afterwards, the membranes were washed with TBS-T, incubated with the according horseradish peroxidase-linked secondary antibody (1:10,000) (HRP linked goat anti-mouse (A4416), HRP linked rabbit anti-goat (A5420), HRP linked goat anti-rabbit (A0545): Sigma-Aldrich, St. Louis, MO), washed again with TBS-T, and incubated with ECL-solution (0.2 mM p-coumaric acid, 1.25 mM luminol, 0.009% H_2_O_2_ in 100 mM Tris-HCl (pH 8.5)) or Western Bright Chemiluminescent substrate solution (541005, Biozym Scientific, Hessisch Oldendorf, Germany). The chemiluminescence signal was detected on X-ray films (PIER34089, VWR, Radnor, PA), quantified using Fiji ImageJ (NIH, Bethesda, MD), and normalized to β-actin.

For the same experiment (same figure panel), immunoblots were performed side by side.

### TG Quantification

Snap-frozen samples of liver and iBAT were homogenized in isopropanol (1 ml per 100 mg tissue) using an ultra-turrax (T10, IKA, Staufen, Germany). The homogenates were incubated for 1h at 60°C and centrifuged at 4°C for 15 min at 3,000 x *g*. The pellet was resuspended in isopropanol, incubated again for 1h at 60°C, and centrifuged at 4°C for 15 min at 3,000 x *g.* Both supernatants were combined and the TG content was measured using a Triglycerides Quantification Kit (183000, Greiner Diagnostics, Bahlingen am Kaiserstuhl, Germany) according to the manufacturer’s instructions.

### Glycogen quantification

Snap-frozen iBAT samples were homogenized in deionized water (200 µl per 10 mg tissue) using a TissueLyser bead mill (85210, Qiagen, Hilden, Germany). The glycogen content of the homogenates was measured using a Glycogen Assay Kit II (ab169558, Abcam, Cambridge, United Kingdom) according to the manufacturer’s instructions.

### Animal monitoring

To measure the metabolic rate of the mice, the animals were studied in a climate controlled indirect calorimetry animal monitoring system (PhenoMaster, TSE Systems GmbH, Bad Homburg, Germany). The mice were adapted to the cages for 48h before the measurement with free access to water and food, and were maintained under 12h light/12h dark cycles either at room temperature (22°C) or exposed to cold (5°C). For each cage, the food intake, water consumption, locomotor activity, CO_2_ production, and O_2_ consumption was measured in 15 min intervals.

### Metabolic tracer study

Mice fasted for 4h received an oral gavage of 250 µl 20% intralipid. The gavaged lipid emulsion contained 47 mg triglycerides / kg body weight and [^14^C]triolein (1 MBq/kg) (ARC0291, Hartmann Analytik, Braunschweig, Germany). Afterwards, the mice were injected intraperitoneally with 0.9% NaCl solution containing [^3^H]deoxyglucose (1 MBq/kg) (MT911, Hartmann Analytik, Braunschweig, Germany). After 1h, the mice were anesthetized and organs were harvested after systemic perfusion with PBS-heparin (10 U/ml) via the left heart ventricle. The tissue samples were dissolved in SolvableTM (6NE9100, Perkin Elmer, Waltham, MA) at 60°C and disintegrations per minute (dpm) were measured by liquid scintillation counting.

### *Ex vivo* [^14^C]-Glucose incorporation assay

Mice housed at 5°C for 48 h were sacrificed, iBAT depots were freed from WAT, and kept in DMEM media (D6429, Sigma-Aldrich, St. Louis, MO). iBAT depots were cut into small pieces to increase surface area. All iBAT pieces from one mouse were transferred into a well of a 12-well dish containing 0.5 ml high-glucose DMEM supplemented with 2% FA-free BSA (A6003, Sigma-Aldrich, St. Louis, MO) and 0.4 µCi [^14^C(U)]-D-glucose (ARC0122, Hartmann Analytik, Braunschweig, Germany). iBAT slices were incubated for 4 h at 37°C in humidified atmosphere and 5% CO_2_. Then, iBAT slices were washed in PBS and lipids were extracted twice using CHCl_3_:methanol (2:1, v/v) supplemented with 0.1% acetic acid. Lipid extracts were pooled and an aliquot of samples and standard mix (triolein:diolein:monoolein:oleic acid) were applied to a silica 60 thin-layer chromatography plate (1.05553.0001, Merck, Darmstadt, Germany) using hexane:diethylether:acetic acid (70:29:1, v/v/v) as mobile phase. Upon chromatographic separation, lipids were visualized by iodine vapor and bands corresponding to triolein and fatty acids were cut out. The co-migrating radioactivity was analyzed by scintillation counting and expressed as cpm/iBAT depot.

### Glucose and insulin tolerance

To analyze whole body glucose tolerance, mice were fasted for 6h and blood glucose was measured using an Accu-Chek® Active blood glucometer (Roche Diagnostics GmbH, Mannheim, Germany). Then, the mice were intraperitoneally injected with a glucose solution (2 g glucose/kg body weight) and blood glucose was determined after 15, 30, 60, 90, and 120 min. For the insulin tolerance test, mice were fasted for 4h and the initial blood glucose was measured. Afterwards, the mice received an intraperitoneal injection of an insulin solution (Actrapid, Novo Nordisk A/S, Bagsvaerd, Denmark) (0.75 insulin units / kg body weight) and blood glucose was determined after 15, 30, 60, 90, and 120 min.

### Plasma membrane enrichment

The plasma membrane fractionation was performed as described 83 with minor modifications. Snap-frozen samples of iBAT were homogenized in NP40-free lysis buffer (0.5 mM DTT in 50 mM Tris-HCl (pH 8.0) and freshly added protease and phosphatase inhibitors) using a tapered tissue grinder (DWK Life Sciences, Milville, New Jersey). The homogenate was passed through a 25G needle 6 times and centrifuged for 10 min at 1,000 x *g* and 4°C (pellet: P1, supernatant: S1). The pellet was resuspended in NP40-free lysis buffer and shaken for 10 min at 4°C. Afterwards, the solution was centrifuged for 10 min at 1,000 x *g* and 4°C (pellet: P2, supernatant: S2). This pellet was resuspended in NP-40 lysis buffer (1% NP-40 in NP40-free lysis buffer; S4) and incubated for 1h at 4°C with shaking. This suspension and the combined supernatants S1 and S2 (combined: S3) were centrifuged for 20 min at 16,000 x *g* and 4°C. The supernatant of S4 was designated as PM fraction and the supernatant of S3 was designated as PM-free fraction. The protein analysis of the two fractions was performed by Western blotting analysis (see above).

### Electron microscopy

Formaldehyde fixed tissue samples were further stabilized by means of high-pressure freezing and freeze-substitution, followed by epoxy resin embedding as previously described 84. Sections of 100 nm were subjected to PAS cytochemistry 85. Alternatively, formaldehyde tissue was cryo-protected with 2.3M sucrose and subjected to cryo-sectioning 84. Sections were imaged by transmission electron microscopy.

### Cell culture

LT2 flox/- and LT2-/- MEFs from Teis et.al.^25^ were used in this study. All cells were cultured in DMEM with high glucose (D6429, Sigma-Aldrich, St. Louis, MO) supplemented with 10% FBS (10270-106, Gibco, Waltham, MA) and 1% penicillin/ streptomycin (P0781, Sigma-Aldrich, St. Louis, MO) and maintained at 37°C, 5% CO_2,_ and 95% humidity. All cell lines were regularly tested negative for mycoplasm infection. Confluent cells were passaged by trypsination using 1x trypsin solution (T4174, Sigma-Aldrich, St. Louis, MO) and phosphate-buffered saline. For the starvation experiment, cells were serum-starved in DMEM with high glucose and penicillin/streptomycin for 5h. Cells were treated with 1 µM ZSTK-474 (HY-50847-10mg, MedChemExpress, South Brunswick, NJ) for 15 min and/or 100 nM insulin (I0516, Sigma-Aldrich, St. Louis, MO) for 10 min. Afterwards, cells were harvested at the indicated time point in ice cold PBS using a rubber policeman. Cells were pelleted at 1,000 x *g* for 5 min and snap-frozen in liquid nitrogen. For Western blotting analysis, the cell pellets were lysed for 30 min at room temperature using a cell lysis buffer (50 mM Tris-HCl pH 8.0, 150 mM NaCl, 1% Triton X-100, 10% glycerol, 1 mM EDTA, 50 mM NaF, 5 mM Na_4_P_2_O_7_, 1 mM Na_3_VO_4_, 0.5 mM PMSF and freshly added protease inhibitors). Protein content was measured using the Bradford Assay Kit (23236, ThermoFisher Scientific, Waltham, MA).

### Crispr-mediated knock-out of Rictor and reconstitution of LAMTOR2

To generate Rictor knockout cells, guide RNAs (gRNAs) targeting *Rictor* were selected using the online tool CHOPCHOP (http://chopchop.cbu.uib.no/) and were cloned into the lentiviral vector pLentiCRISPRv2-LoxPv1 (based on the pLentiCRISPRv2 plasmid from Addgene 5261 with inserted loxP sites flanking the elongation factor 1α short promoter). To generate lentiviral particles containing the CRISPR/Cas9 and gRNA encoding vectors, HEK293T cells were transfected using PEI (23966-1, Polyscience, Warrington, PA) with the pLentiCRISPRv2-LoxPv1 gRNA containing plasmid, pVSV-G (631530, Clontech, Mountain View, CA) and psPAX2. The virus particle containing medium supernatant was used to infect LAMTOR2-/- MEFs in presence of polybrene. These MEFs were selected with 1 µg/ml puromycin and single cell clones were grown and tested by PCR using the respective primers and Western Blot analysis (see above).

For reconstituting LT2 in LT2-/- MEFs and LT2-/- Rictor-/- MEFs, an already constructed LT2 flanked by HA and Strep-tag containing pENTR4 vector was used to generate a pCCL-HA-LT2 vector, in which the LAMTOR2 construct is under the control of an EF1alpha promoter. The pCCL-HA-LT2 contains a blasticidin resistance gene. Lentiviral particles containing the pCCL vector were generated in HEK293T cells by transfecting the HEK293T cells with the pCCL vector and pVSV-G and psPAX2 (see above).

LT2-/- MEFs and LT2-/- Rictor-/- MEFs were infected by the lentivirus containing medium in presence of polybrene and selected with 0.6 µg/ml blasticidin. Single cell clones were grown and tested for HA- LT2 expression by PCR and Western Blot analysis.

### Quantification and statistical analysis

All data presented are biological replicates and bar charts are represent mean±SD with single values as dots. Unpaired two-tailed Student’s t-tests were used for comparing two groups. For more than two groups, two-way ANOVA and multiple comparison correction for p-values was used. The statistical parameters are specified in the figure legends and the data was analyzed with GraphPad Prism. No method was used to check for exclusion of values. As test for normality distribution, we checked the data distribution in Q-Q Plots. Differences were considered statistically significant for p-values below 0.05 and p-values are depicted in the charts.

For the animal experiments, sample sizes were determined using a power analysis based on a 50% difference, a power of 90% and a variance depending on previous animal experiments. Nevertheless, in consent with “3R” rule a lower sample sizes were used whenever possible . For cell culture experiments, at least 4 biological replicates were performed according to previous experiments in our laboratory.

No blinding or randomization of the samples were applied.

## Acknowledgments

This work was supported by the Austrian Science Fund FWF (PhD program MCBO (W 1101) to GLi, P26682 to LAH, W1226, F73 and P30882 to DK). L.S. and J.H. were supported by the Deutsche Forschungsgemeinschaft DFG (SCHE522/4-1 and HE3645/10-1). We acknowledge the excellent technical assistance of Johannes Rainer, Karin Gutleben, and Barbara Witting.

## Author Contributions

G.Li., N.V., R.S., M.H., C.K., M.D-M., G.L., M.W.H., T.O.E., S.H. performed the experiments. C.H.S., L.S., J.H. and D.K. designed experiments and contributed conceptually. L.A.H. initiated and supervised the study and together with G.Li. coordinated the study and wrote the manuscript. All authors reviewed and edited the manuscript.

## Competing Interests

The authors declare no competing interests.

## Supplemental Information

**Figure S1.**
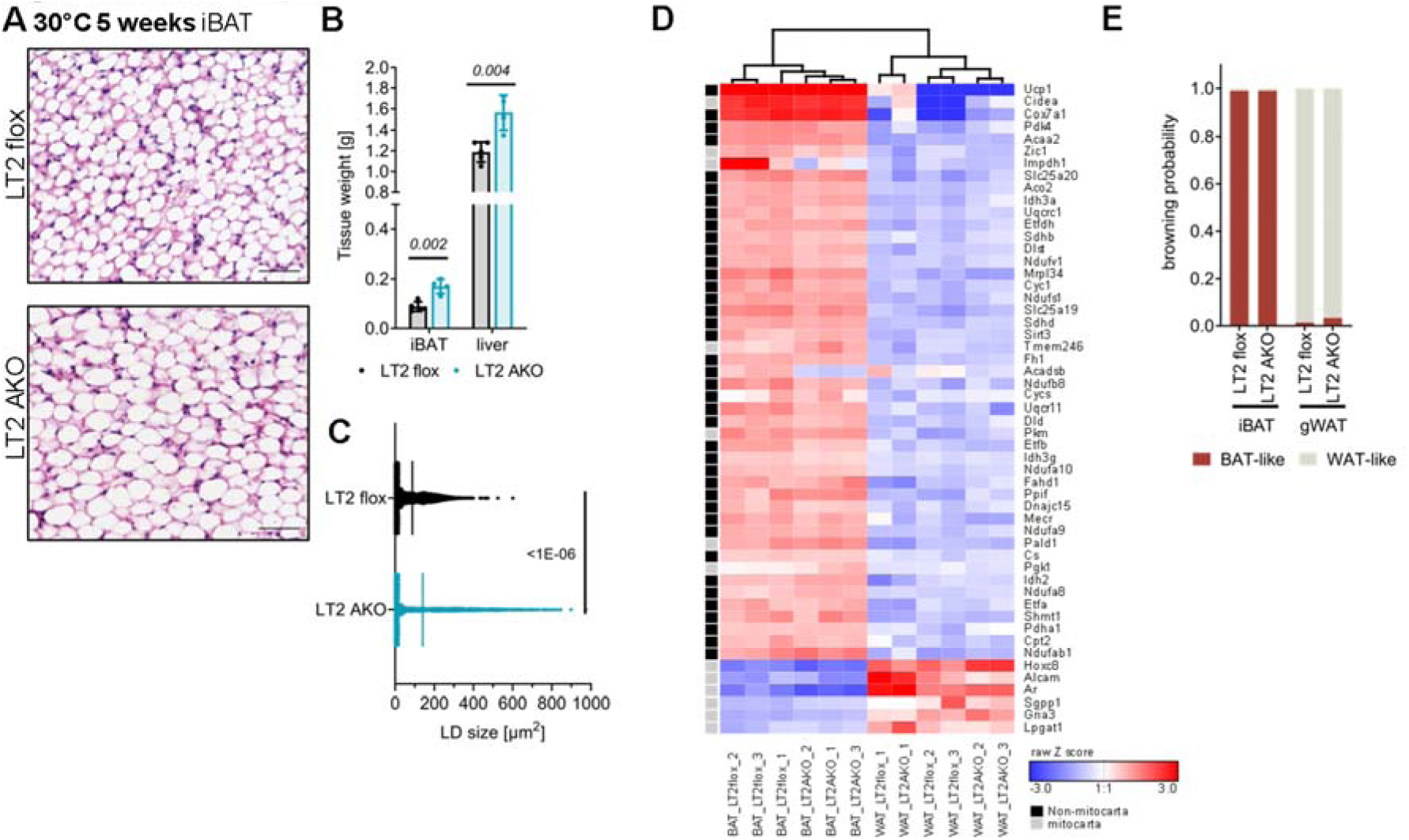
iBAT lipid accumulation in thermoneutral housing and unchanged iBAT marker genes in LT2 AKO mice. (A) Representative H&E staining of iBAT sections of mice acclimatized for 5 weeks at thermoneutrality (scale bar 50 µm) with (B) quantification of iBAT and liver weight as well as (C) LD size (n=5). Microarray analysis was performed using mRNA from iBAT of male mice aged 8-9 weeks. (D) ProFAT analysis of microarray data of iBAT and gWAT from LT2 AKO and LT2 flox mice (n=3). (E) Calculation of the browning probability after ProFAT analysis. Data are presented as mean (n=3). Bar charts present means ± SD with individual values presented as dots. Statistical significance was calculated using two-tailed unpaired Student’s t-tests with italic numbers representing the p-value.

**Figure S2.**
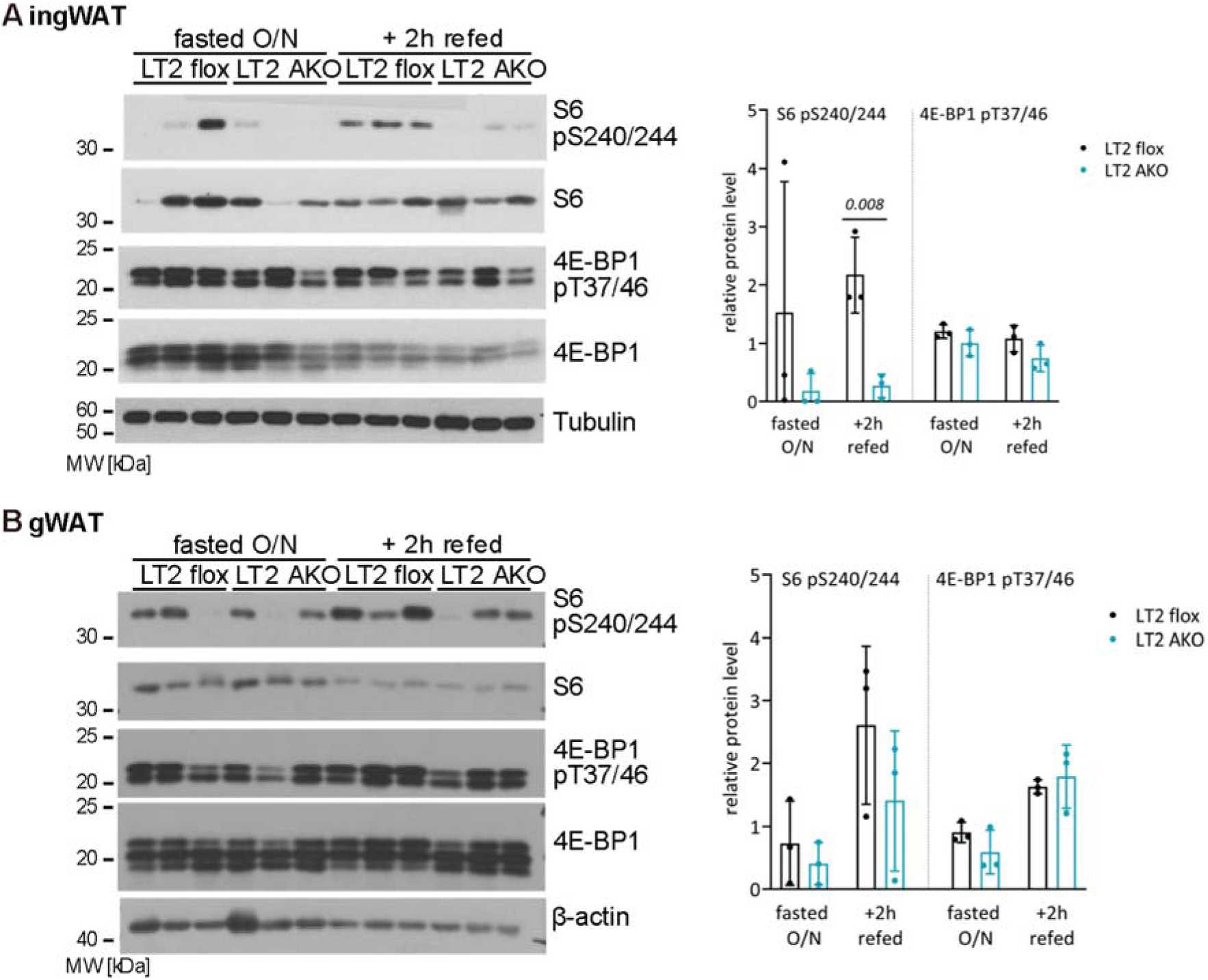
Partially reduced mTORC1 signaling in LT2 AKO ingWAT and inconclusive mTORC1 signaling in LT2 AKO gWAT. Immunoblot and quantification of mTORC1 downstream targets S6 and 4E-BP1 with their respective phosphorylations in (A) ingWAT and (B) gWAT samples of female LT2 flox and LT2 AKO mice (n=3). Bar charts present means ± SD with individual values presented as dots. Statistical significance was calculated using two-tailed unpaired Student’s t-tests with italic numbers representing the p-value.

**Figure S3.**
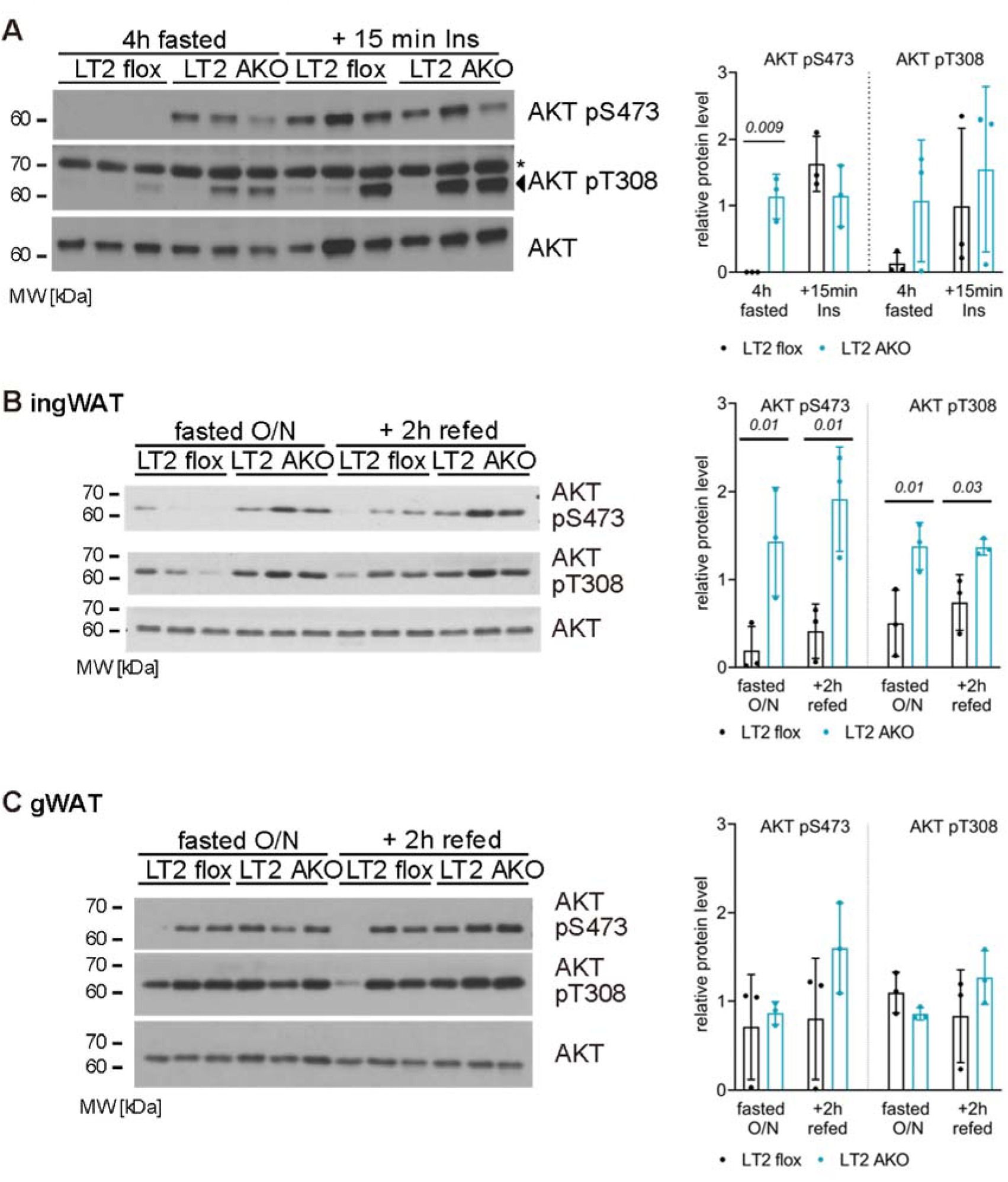
Insulin treatment cannot further increase AKT phosphorylation in LT2 AKO iBAT and increased AKT phosphorylation in LT2 AKO ingWAT, but not in LT2 AKO gWAT. (A) Immunoblots and quantification of AKT and its phosphorylation sites (pS473, pT308) in iBAT lysates of 4h-fasted male mice with or without insulin treatment for 15 min (n=3). Immunoblot and quantification of AKT and its phosphorylation sites (pS473, pT308) in (B) ingWAT and (C) gWAT samples of female LT2 flox and LT2 AKO mice (n=3). Statistical significance was calculated using two-tailed unpaired Student’s t-tests with the italic numbers representing the p-value.

**Figure S4.**
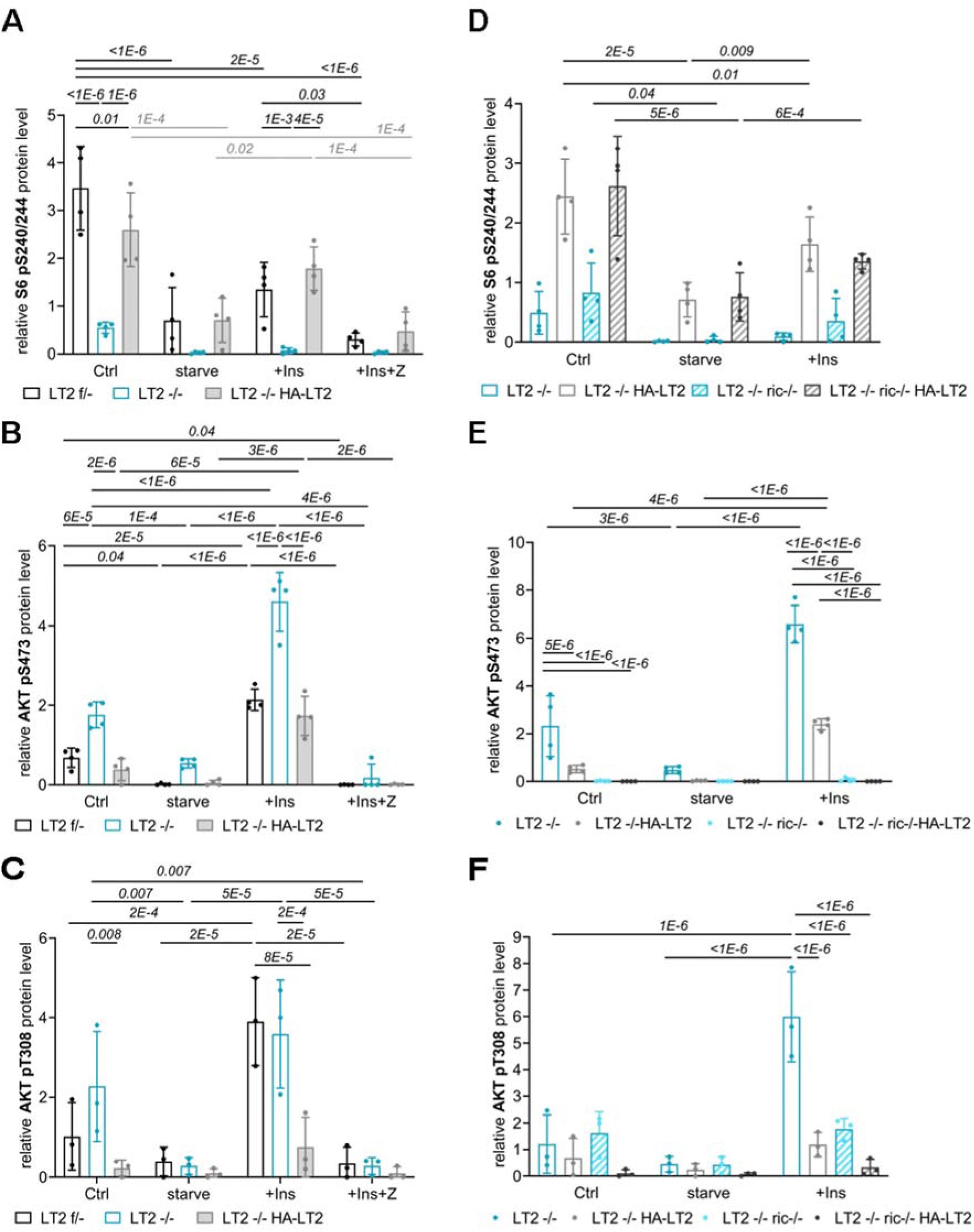
Quantifications of immunoblots from Fig 5. Statistical significance was calculated using two-tailed unpaired Student’s t-tests with multiple comparison correction with italic numbers representing the p-value.

**Figure S5.**
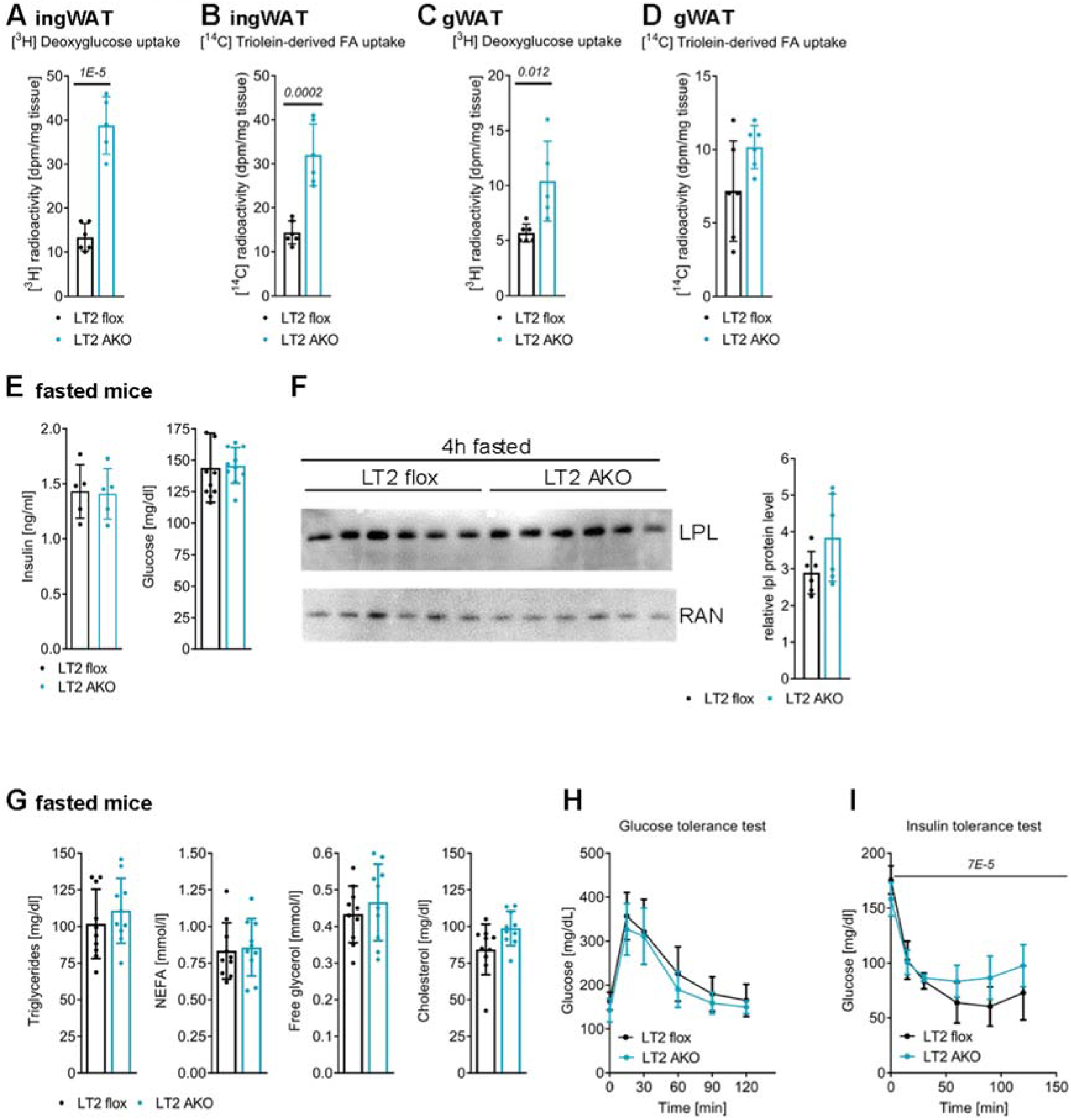
LT2 AKO ingWAT and gWAT have increased glucose uptake, but only ingWAT also increased FA uptake. Overall, LT2 AKO mice are insulin resistant despite unchanged plasma lipid and glucose concentrations. Male mice aged 12-16 weeks were analyzed. Mice were fasted for 4h before they were (A, C) i.p. injected with [^3^H]deoxyglucose (B, D) gavaged with intralipid emulsion containing [^14^C]triolein and and uptake of radioactivity was determined in (A, B) ingWAT and (C, D) gWAT (n=6).(E) Plasma insulin and glucose levels of 4h- fasted mice (n=5). (F) LPL immunoblot and quantification in iBAT lysates from 4h-fasted mice (n=5). (G) Plasma triglyceride, non-esterified fatty acid (NEFA), free glycerol, and cholesterol concentrations of 4h-fasted mice (n=5). (H) Glucose and (I) insulin tolerance test in 6h- or 4h-fasted mice, respectively (n=5). Bar charts present means ± SD with individual values presented as dots. Statistical significance was calculated using (A - G) two-tailed unpaired Student’s t-tests or (H, I) ANOVA analysis with the italic number representing the p-value.

**Figure S6.**
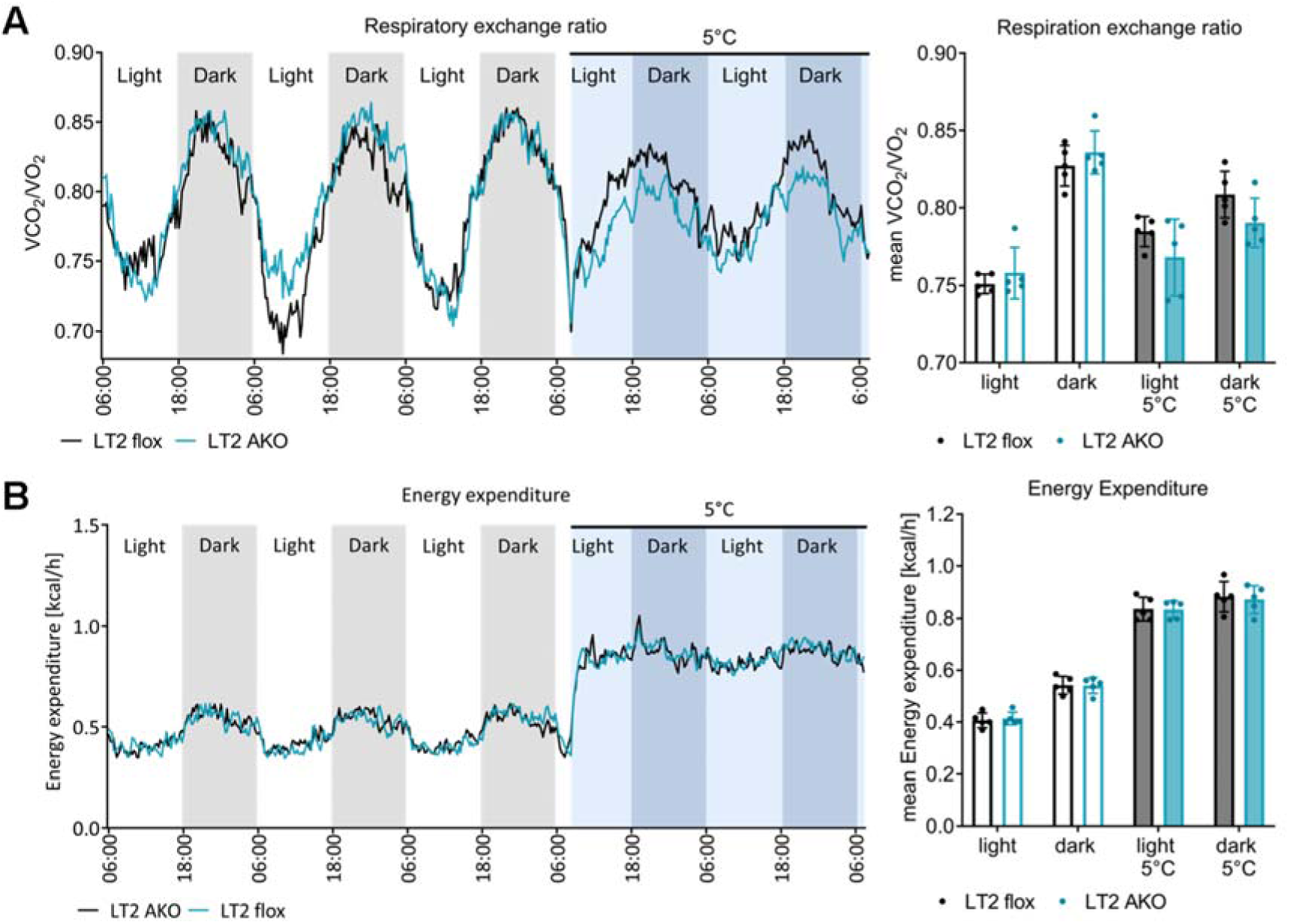
Unchanged *in vivo* metabolic parameters of LT2 AKO housed at 5°C for 48h. Male mice (30 weeks old) were housed at 22°C before they were exposed to 5°C for 48h (n=5). (A) Respiratory exchange ratio and (B) energy expenditure over 72h at room temperature and 48h at 5°C. Bar charts present means ± SD with individual values presented as dots. Statistical significance was calculated using two-tailed unpaired Student’s t-tests.

**Table S1.**
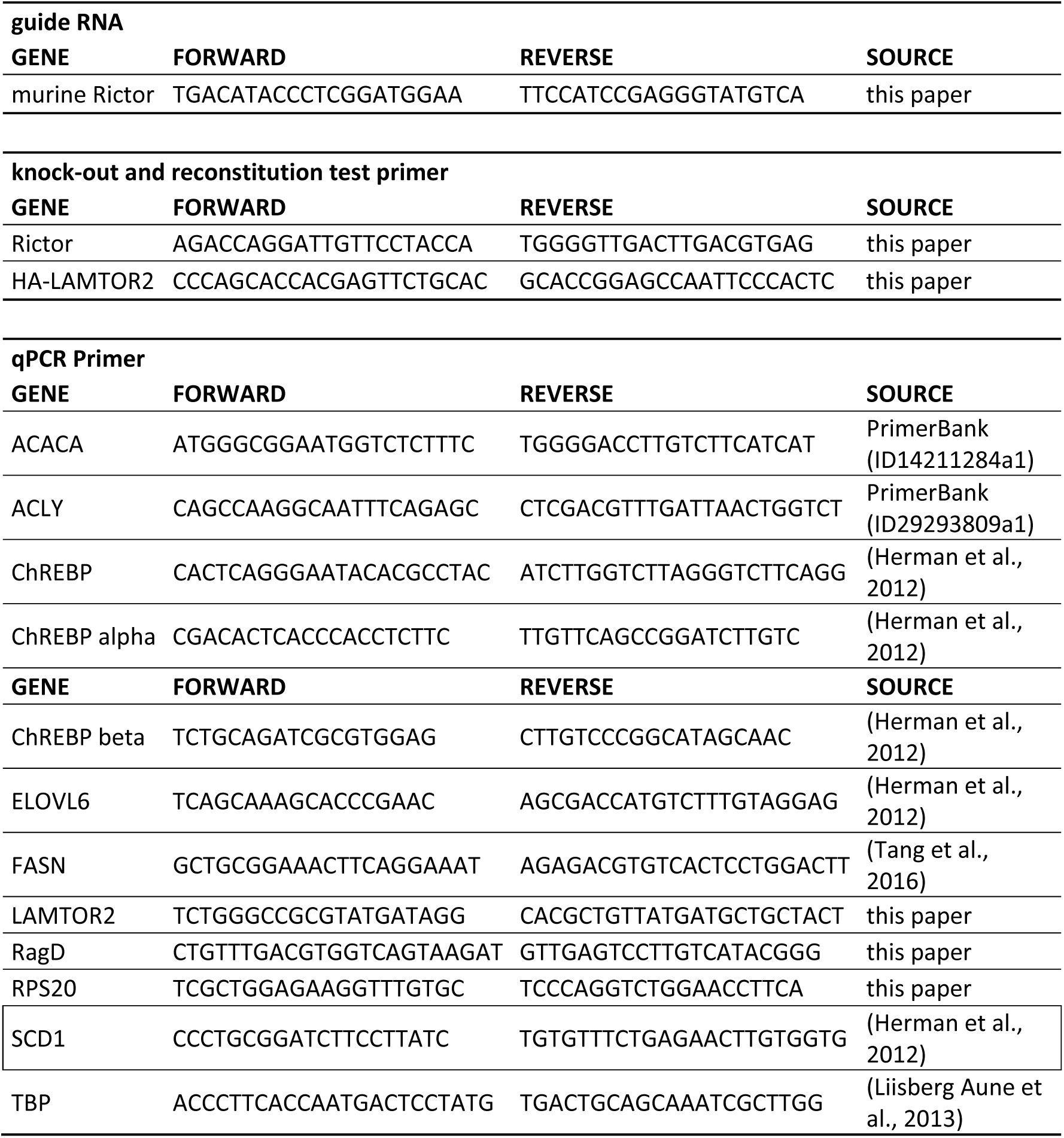
Primers used as gRNA for the generation of the Rictor -/- MEFs, for testing the deletion of Rictor and expression of HA-LT2 in MEFs as well as primers for relative gene expression analysis.

## Notes

### Competing Interest Statement

The authors have declared no competing interest.

## References

1. Cannon, B. & Nedergaard, J. Brown Adipose Tissue: Function and Physiological Significance. Physiological Reviews 84, 277–359 (2004).

2. Townsend, K. L. & Tseng, Y. H. Brown fat fuel utilization and thermogenesis. Trends in Endocrinology and Metabolism 25, 168–177 (2014).

3. Aquila, H., Link, T. A. & Klingenberg, M. The uncoupling protein from brown fat mitochondria is related to the mitochondrial ADP/ATP carrier. Analysis of sequence homologies and of folding of the protein in the membrane. The EMBO journal 4, 2369–2376 (1985).

4. Heaton, G. M., Wagenvoord, R. J., Kemp, A. & Nicholls, D. G. Brown-adipose-tissue mitochondria: photoaffinity labelling of the regulatory site of energy dissipation. European journal of biochemistry 82, 515–21 (1978).

5. Schreiber, R. et al. Cold-Induced Thermogenesis Depends on ATGL-Mediated Lipolysis in Cardiac Muscle, but Not Brown Adipose Tissue. Cell Metabolism 26, 753–763.e7 (2017).

6. Shin, H. et al. Lipolysis in Brown Adipocytes Is Not Essential for Cold-Induced Thermogenesis in Mice. Cell Metabolism 26, 764–777.e5 (2017).

7. Bartelt, A. et al. Brown adipose tissue activity controls triglyceride clearance. Nature Medicine 17, 200–206 (2011).

8. Fischer, A. W. et al. Lysosomal lipoprotein processing in endothelial cells stimulates adipose tissue thermogenic adaptation. Cell Metabolism 33, 547–564.e7 (2021).

9. Heine, M. et al. Lipolysis Triggers a Systemic Insulin Response Essential for Efficient Energy Replenishment of Activated Brown Adipose Tissue in Mice. Cell Metabolism 28, 644–655.e4 (2018).

10. Knudsen, J. R. et al. Growth Factor-Dependent and -Independent Activation of mTORC2. Trends in Endocrinology and Metabolism 31, 13–24 (2020).

11. Lee, P. L., Jung, S. M. & Guertin, D. A. The Complex Roles of Mechanistic Target of Rapamycin in Adipocytes and Beyond. Trends in Endocrinology and Metabolism 28, 319–339 (2017).

12. Petersen, M. C. & Shulman, G. I. Mechanisms of insulin action and insulin resistance. Physiological Reviews 98, 2133–2223 (2018).

13. Larance, M. et al. Characterization of the role of the Rab GTPase-activating protein AS160 in insulin-regulated GLUT4 trafficking. Journal of Biological Chemistry 280, 37803–37813 (2005).

14. Menon, S. et al. Spatial control of the TSC complex integrates insulin and nutrient regulation of mTORC1 at the lysosome. Cell 156, 771–785 (2014).

15. Bar-Peled, L., Schweitzer, L. D., Zoncu, R. & Sabatini, D. M. Ragulator is a GEF for the rag GTPases that signal amino acid levels to mTORC1. Cell 150, 1196–1208 (2012).

16. Sancak, Y. et al. The rag GTPases bind raptor and mediate amino acid signaling to mTORC1. Science 320, 1496–1501 (2008).

17. Sancak, Y. et al. Ragulator-rag complex targets mTORC1 to the lysosomal surface and is necessary for its activation by amino acids. Cell 141, 290–303 (2010).

18. Labbé, S. M. et al. mTORC1 is Required for Brown Adipose Tissue Recruitment and Metabolic Adaptation to Cold. Scientific reports 6, 37223 (2016).

19. Lee, P. L., Tang, Y., Li, H. & Guertin, D. A. Raptor/mTORC1 loss in adipocytes causes progressive lipodystrophy and fatty liver disease. Molecular Metabolism 5, 422–432 (2016).

20. Shan, T. et al. Adipocyte-specific deletion of mTOR inhibits adipose tissue development and causes insulin resistance in mice. Diabetologia 59, 1995–2004 (2016).

21. Wada, S. et al. The tumor suppressor FLCN mediates an alternate mTOR pathway to regulate browning of adipose tissue. Genes and Development 30, 2551–2564 (2016).

22. Peterson, T. R. et al. MTOR complex 1 regulates lipin 1 localization to control the srebp pathway. Cell 146, 408–420 (2011).

23. Yecies, J. L. et al. Akt stimulates hepatic SREBP1c and lipogenesis through parallel mTORC1- dependent and independent pathways. Cell Metabolism 14, 21–32 (2011).

24. Nada, S. et al. The novel lipid raft adaptor p18 controls endosome dynamics by anchoring the MEK-ERK pathway to late endosomes. EMBO Journal 28, 477–489 (2009).

25. Teis, D., Wunderlich, W. & Huber, L. A. Localization of the MP1-MAPK scaffold complex to endosomes is mediated by p14 and required for signal transduction. Developmental Cell 3, 803– 814 (2002).

26. Zhang, C. S. et al. The lysosomal v-ATPase-ragulator complex is a common activator for AMPK and mTORC1, acting as a switch between catabolism and anabolism. Cell Metabolism 20, 526– 540 (2014).

27. De Araújo, M. E. G. et al. Stability of the endosomal scaffold protein lAMTOR3 depends on heterodimer assembly and proteasomal degradation. Journal of Biological Chemistry 288, 18228–18242 (2013).

28. Harrington, L. S. et al. The TSC1-2 tumor suppressor controls insulin-PI3K signaling via regulation of IRS proteins. Journal of Cell Biology 166, 213–223 (2004).

29. Tremblay, F. et al. Identification of IRS-1 Ser-1101 as a target of S6K1 in nutrient- and obesity- induced insulin resistance. Proceedings of the National Academy of Sciences of the United States of America 104, 14056–14061 (2007).

30. Wang, L. et al. Peripheral Disruption of the Grb10 Gene Enhances Insulin Signaling and Sensitivity In Vivo. Molecular and Cellular Biology 27, 6497–6505 (2007).

31. Hsu, P. P. et al. The mTOR-regulated phosphoproteome reveals a mechanism of mTORC1- mediated inhibition of growth factor signaling. Science 332, 1317–1322 (2011).

32. Yu, Y. et al. Phosphoproteomic analysis identifies Grb10 as an mTORC1 substrate that negatively regulates insulin signaling. Science 332, 1322–1326 (2011).

33. Cheng, Y. et al. Prediction of Adipose Browning Capacity by Systematic Integration of Transcriptional Profiles. Cell Reports 23, 3112–3125 (2018).

34. Yoon, M. S. The role of mammalian target of rapamycin (mTOR) in insulin signaling. Nutrients 9, (2017).

35. Zhang, R., Lahens, N. F., Ballance, H. I., Hughes, M. E. & Hogenesch, J. B. A circadian gene expression atlas in mammals: Implications for biology and medicine. Proceedings of the National Academy of Sciences of the United States of America 111, 16219–16224 (2014).

36. Baar, E. L., Carbajal, K. A., Ong, I. M. & Lamming, D. W. Sex- and tissue-specific changes in mTOR signaling with age in C57BL/6J mice. Aging Cell 15, 155–166 (2016).

37. Demirkan, G., Yu, K., Boylan, J. M., Salomon, A. R. & Gruppuso, P. A. Phosphoproteomic profiling of in vivo signaling in liver by the mammalian target of rapamycin complex 1 (mTORC1). PloS one 6, e21729 (2011).

38. Settembre, C. et al. A lysosome-to-nucleus signalling mechanism senses and regulates the lysosome via mTOR and TFEB. EMBO Journal 31, 1095–1108 (2012).

39. Martina, J. A., Chen, Y., Gucek, M. & Puertollano, R. MTORC1 functions as a transcriptional regulator of autophagy by preventing nuclear transport of TFEB. Autophagy 8, 903–914 (2012).

40. Di Malta, C. et al. Transcriptional activation of RagD GTPase controls mTORC1 and promotes cancer growth. Science 356, 1188–1193 (2017).

41. Sass, F. et al. TFEB deficiency attenuates mitochondrial degradation upon brown adipose tissue whitening at thermoneutrality. Molecular Metabolism 47, 101173 (2021).

42. Yaguchi, S. I. et al. Antitumor activity of ZSTK474, a new phosphatidylinositol 3-kinase inhibitor. Journal of the National Cancer Institute 98, 545–556 (2006).

43. Lobo, S. & Bernlohr, D. A. Fatty acid transport in adipocytes and the development of insulin resistance. Novartis Foundation Symposium 286, 113–121 (2007).

44. Geraghty, K. M. et al. Regulation of multisite phosphorylation and 14-3-3 binding of AS160 in response to IGF-1, EGF, PMA and AICAR. Biochemical Journal 407, 231–241 (2007).

45. Kane, S. et al. A method to identify serine kinase substrates. Akt phosphorylates a novel adipocyte protein with a Rab GTPase-activating protein (GAP) domain. Journal of Biological Chemistry 277, 22115–22118 (2002).

46. Sano, H. et al. Insulin-stimulated phosphorylation of a Rab GTPase-activating protein regulates GLUT4 translocation. Journal of Biological Chemistry 278, 14599–14602 (2003).

47. Herman, M. A. et al. A novel ChREBP isoform in adipose tissue regulates systemic glucose metabolism. Nature 484, 333–338 (2012).

48. Ishii, S., Iizuka, K., Miller, B. C. & Uyeda, K. Carbohydrate response element binding protein directly promotes lipogenic enzyme gene transcription. Proceedings of the National Academy of Sciences of the United States of America 101, 15597–15602 (2004).

49. Schlein, C. et al. Endogenous Fatty Acid Synthesis Drives Brown Adipose Tissue Involution. Cell reports 34, 108624 (2021).

50. Sanchez-Gurmaches, J. et al. Brown Fat AKT2 Is a Cold-Induced Kinase that Stimulates ChREBP-Mediated De Novo Lipogenesis to Optimize Fuel Storage and Thermogenesis. Cell Metabolism 27, 195–209.e6 (2018).

51. Guan, D. et al. Diet-Induced Circadian Enhancer Remodeling Synchronizes Opposing Hepatic Lipid Metabolic Processes. Cell 174, 831–842.e12 (2018).

52. Horton, J. D., Goldstein, J. L. & Brown, M. S. SREBPs: activators of the complete program of cholesterol and fatty acid synthesis in the liver. Journal of Clinical Investigation 109, 1125–1131 (2002).

53. Goldstein, J. L., DeBose-Boyd, R. A. & Brown, M. S. Protein sensors for membrane sterols. Cell 124, 35–46 (2006).

54. Morigny, P., Boucher, J., Arner, P. & Langin, D. Lipid and glucose metabolism in white adipocytes: pathways, dysfunction and therapeutics. Nature Reviews Endocrinology 17, 276– 295 (2021).

55. Polak, P. et al. Adipose-Specific Knockout of raptor Results in Lean Mice with Enhanced Mitochondrial Respiration. Cell Metabolism 8, 399–410 (2008).

56. Brown, M. S. & Goldstein, J. L. Cholesterol feedback: From Schoenheimer’s bottle to Scap’s MELADL. Journal of Lipid Research 50, 15–28 (2009).

57. Scheffler, J. M. et al. LAMTOR2 regulates dendritic cell homeostasis through FLT3-dependent mTOR signalling. Nature communications 5, 5138 (2014).

58. Carmean, C. M., Bobe, A. M., Yu, J. C., Volden, P. A. & Brady, M. J. Refeeding-induced brown adipose tissue glycogen hyper-accumulation in mice is mediated by insulin and catecholamines. PloS one 8, e67807 (2013).

59. Farkas, V., Kelenyi, G. & Sandor, A. A dramatic accumulation of glycogen in the brown adipose tissue of rats following recovery from cold exposure. Archives of biochemistry and biophysics 365, 54–61 (1999).

60. Rahman, S. M. et al. Stearoyl-CoA desaturase 1 deficiency increases insulin signaling and glycogen accumulation in brown adipose tissue. American Journal of Physiology - Endocrinology and Metabolism 288, E381–E387 (2005).

61. Chitraju, C., Fischer, A. W., Farese, R. v. & Walther, T. C. Lipid Droplets in Brown Adipose Tissue Are Dispensable for Cold-Induced Thermogenesis. Cell reports 33, 108348 (2020).

62. Keinan, O. et al. Glycogen metabolism links glucose homeostasis to thermogenesis in adipocytes. Nature 599, (2021).

63. Albert, V. et al. mTORC 2 sustains thermogenesis via Akt-induced glucose uptake and glycolysis in brown adipose tissue . EMBO Molecular Medicine 8, 232–246 (2016).

64. Hung, C. M. et al. Rictor/mTORC2 loss in the Myf5 lineage reprograms brown fat metabolism and protects mice against obesity and metabolic disease. Cell Reports 8, 256–271 (2014).

65. Shearin, A. L., Monks, B. R., Seale, P. & Birnbaum, M. J. Lack of AKT in adipocytes causes severe lipodystrophy. Molecular Metabolism 5, 472–479 (2016).

66. Vogel, G. F. et al. Ultrastructural Morphometry Points to a New Role for LAMTOR2 in Regulating the Endo/Lysosomal System. Traffic 16, 617–634 (2015).

67. Braccini, L. et al. PI3K-C2γ is a Rab5 effector selectively controlling endosomal Akt2 activation downstream of insulin signalling. Nature Communications 6, 7400 (2015).

68. Franke, T. F., Kaplan, D. R., Cantley, L. C. & Toker, A. Direct regulation of the Akt proto- oncogene product by phosphatidylinositol-3,4-bisphosphate. Science 275, 665–668 (1997).

69. Jia, R. & Bonifacino, J. S. Lysosome Positioning Influences mTORC2 and AKT Signaling. Molecular Cell 75, 26–38.e3 (2019).

70. Filipek, P. A. et al. LAMTOR/Ragulator is a negative regulator of Arl8b- and BORC-dependent late endosomal positioning. The Journal of cell biology 216, 4199–4215 (2017).

71. Pu, J. et al. BORC, a Multisubunit Complex that Regulates Lysosome Positioning. Developmental Cell 33, 176–188 (2015).

72. Pu, J., Keren-Kaplan, T. & Bonifacino, J. S. A Ragulator-BORC interaction controls lysosome positioning in response to amino acid availability. Journal of Cell Biology 216, 4183–4197 (2017).

73. Teis, D. et al. p14-MP1-MEK1 signaling regulates endosomal traffic and cellular proliferation during tissue homeostasis. Journal of Cell Biology 175, 861–868 (2006).

74. Eguchi, J. et al. Transcriptional control of adipose lipid handling by IRF4. Cell Metabolism 13, 249–259 (2011).

75. Schmidt, S. et al. Identification of glucocorticoid-response genes in children with acute lymphoblastic leukemia. Blood 107, 2061–2069 (2006).

76. Gentleman, R. C. et al. Bioconductor: open software development for computational biology and bioinformatics. Genome biology 5, R80 (2004).

77. Wu, Z., Irizarry, R. A., Gentleman, R., Martinez-Murillo, F. & Spencer, F. A model-based background adjustment for oligonucleotide expression arrays. Journal of the American Statistical Association 99, 909–917 (2004).

78. Smyth, G. K. Linear models and empirical bayes methods for assessing differential expression in microarray experiments. Statistical applications in genetics and molecular biology 3, Article3 (2004).

79. Benjamini, Y. & Hochberg, Y. Controlling the False Discovery Rate: A Practical and Powerful Approach to Multiple Testing. Journal of the Royal Statistical Society: Series B (Methodological) 57, 289–300 (1995).

80. Xie, Z. et al. Gene Set Knowledge Discovery with Enrichr. Current Protocols 1, 1–51 (2021).

81. Kuleshov, M. v, et al. Enrichr: a comprehensive gene set enrichment analysis web server 2016 update. Nucleic acids research 44, W90–W97 (2016).

82. Chen, E. Y. et al. Enrichr: interactive and collaborative HTML5 gene list enrichment analysis tool. BMC bioinformatics 14, 128 (2013).

83. Yamamoto, N., Yamashita, Y., Yoshioka, Y., Nishiumi, S. & Ashida, H. Rapid Preparation of a Plasma Membrane Fraction: Western Blot Detection of Translocated Glucose Transporter 4 from Plasma Membrane of Muscle and Adipose Cells and Tissues. Current protocols in protein science 85, 29.18.1–29.18.12 (2016).

84. Vogel, G. F. et al. Abnormal Rab11-Rab8-vesicles cluster in enterocytes of patients with microvillus inclusion disease. Traffic 18, 453–464 (2017).

85. Thiery, JP. Mise en evidence des polysaccharides sur coupes fines en microscopie electronique. J Microsc 6, 987–1018 (1967).

